# In vivo serial passaging of human-simian immunodeficiency virus clones identifies viral characteristics that enhance persistent viral replication

**DOI:** 10.1101/2021.09.15.460387

**Authors:** Rajesh Thippeshappa, Patricia Polacino, Shaswath S. Chandrasekar, Khanghy Truong, Anisha Misra, Shiu-Lok Hu, Deepak Kaushal, Jason T. Kimata

## Abstract

We previously reported that a human immunodeficiency virus type 1 with a simian immunodeficiency virus *vif* substitution (HSIV-vifNL4-3) could replicate in pigtailed macaques (PTMs), demonstrating that Vif is a species-specific tropism factor of primate lentiviruses. However, infections did not result in high peak viremia or setpoint plasma viral loads, as observed during SIV infection of PTMs. Here, we characterized variants isolated from one of the original infected animals with CD4 depletion after nearly four years of infection to identify determinants of increased replication fitness. In our studies, we found that the HSIV-vif clones did not express the HIV-1 Vpr protein due to interference from the *vpx* open reading frame in singly spliced *vpr* mRNA. To examine whether these viral genes contribute to persistent viral replication, we generated infectious HSIV-vif clones expressing either the HIV-1 Vpr or SIV Vpx protein. And then to determine viral fitness determinants of HSIV-vif, we conducted three rounds of serial in vivo passaging in PTMs, starting with an initial inoculum containing a mixture of CXCR4-tropic (Vpr^-^ HSIV-vif_NL4-3_ isolated at 196 (C/196) and 200 (C/200) weeks post-infection from a PTM with depressed CD4 counts) and CCR5-tropic HSIV (Vpr^+^ HSIV-vif derivatives based NL-AD8 and Bru-Yu2 and a Vpx expressing HSIV-vif_Yu2_). Interestingly, all infected PTMs showed peak plasma viremia close to or above 10^5^ copies/ml and persistent viral replication for more than 20 weeks. The passage 3 PTM showed peak viral loads greater than 10^5^ viral RNA copies/ml. Infectious molecular clones (IMCs) recovered from the passage 3 PTM (HSIV-P3 IMCs) included mutations required for HIV-1 Vpr expression and those mutations encoded by the CXCR4-tropic HSIV-vif_NL4-3_ isolates C/196 and C/200. The data indicate that the biological isolates selected during long-term infection acquired HIV-1 Vpr expression to enhance their replication fitness in PTMs. Further passaging of HSIV-P3 IMCs in vivo may generate pathogenic variants with higher replication capacity, which will be a valuable resource as challenge virus in vaccine and cure studies.

## Introduction

Several alternate animal models such as infection of macaques with simian immunodeficiency viruses (SIVs) or chimeric simian-human immunodeficiency viruses (SHIVs) have been developed to understand HIV pathogenesis, disease progression, and determine the efficacy of vaccines and drugs. However, the genetic difference between HIV-1 and SIV, and the absence of other HIV-1 genes such as *gag, vif, vpr*, and *nef* in SHIV limits the utility of these models. Therefore, there is a need to rationally and minimally modify HIV-1 such that it can replicate and cause AIDS in macaques. Such an animal model will be a valuable tool for preclinical evaluation of vaccines and the development of novel therapeutic strategies targeting HIV-1 proteins, and for understanding viral immunopathogenesis.

The important lentiviral restriction factors in macaque species such as rhesus macaques (RMs) are the apolipoprotein B mRNA editing enzyme catalytic polypeptide 3 (APOBEC3 or A3) family of proteins, tripartite motif containing (TRIM) family of proteins, BST2/CD317/Tetherin, and sterile alpha motif (SAM) and histidine/aspartic acid (HD) domain containing protein 1 (SAMHD1) (reviewed in (Thippeshappa et al., 2012; Saito and Akari, 2013). However, SIV can overcome RM TRIM5α and the APOBEC3 family of restriction factors and simian-tropic HIV-1 (stHIV-1) or macaque-tropic HIV-1 (mtHIV-1) have been developed by incorporating *capsid* and *vif* sequences from SIVmac239 (Hatziioannou et al., 2006; Saito et al., 2011; Doi et al., 2013; Nomaguchi et al., 2013; Otsuki et al., 2014; Doi et al., 2018). Instead of a full length capsid substitution, an HIV-1 derivative carrying only a short 21 nucleotide segment from the SIV capsid sequence corresponding to the HIV-1 cylophilin A binding loop has also been constructed (Kamada et al., 2006). Additionally, variants with CCR5-tropic HIV-1 have also been developed (Otsuki et al., 2014; Doi et al., 2017). These variants of stHIV-1 or mtHIV-1 establish infection in vivo in different species of nonhuman primates (NHPs) (Igarashi et al., 2007; Saito et al., 2011; Saito et al., 2013; Otsuki et al., 2014; Doi et al., 2018). However, none of the variants result in CD4 depletion, and there remains a need to develop a pathogenic macaque-tropic HIV-1 (reviewed in (Thippeshappa et al., 2020).

Compared to other NHPs used in AIDS research, PTMs are relatively more susceptible to HIV-1 infection (Agy et al., 1992; Frumkin et al., 1993; Gartner et al., 1994a; Gartner et al., 1994b; Agy et al., 1997; Bosch et al., 1997; Bosch et al., 2000). While PTMs can be infected with HIV-1, viral loads diminished rapidly (Agy et al., 1992). Attempts to *in vivo* passage HIV-1 in PTMs failed to select variants capable of persistent replication.

An explanation for the susceptibility of PTM CD4^+^ T cells to HIV-1 is that PTMs do not express restriction factor TRIM5α. Instead, they express novel isoforms of TRIM5 (TRIM5θ and TRIM5η) and TRIM5-cyclophilin A fusion protein (TRIMcyp) that do not interfere with HIV-1 infection (Liao et al., 2007; Brennan et al., 2008; Newman et al., 2008; Virgen et al., 2008). Absence of TRIM5α suggests that other retroviral restriction factors in PTMs such as APOBEC3 family of proteins, BST2, and SAMHD1 may limit replication of HIV-1. Since APOBEC3 family proteins can be degraded by SIVmac and HIV-2 *vif*, Hatziioannou *et al*. constructed minimally modified HIV-1 derivatives carrying either SIVmac *vif* or HIV-2 *vif* (Hatziioannou et al., 2009). PTMs infected intravenously (IV) with a mixture of these two viruses exhibited acute infection and persistent viremia for up to 25 weeks post-infection (wpi). However, CD4^+^ T-cell depletion was not observed in the animals. To select a variant with increased fitness, serial in vivo passaging of a mixture of four clonal HIV-1_NL4-3_–derived viruses, each encoding CCR5-tropic gp120 env from YU2, BaL, AD8, and KB9 was conducted in PTMs transiently depleted of CD8 T cells. Viral swarm or IMC generated following passaging caused CD4 depletion only in macaques that were transiently depleted of CD8 T cells. However, they were controlled in immunocompetent PTMs (Hatziioannou et al., 2014; Schmidt et al., 2019). Inability of passaged viruses to cause AIDS in non CD8-depleted macaques suggest partial adaptation to the PTM host. Despite these studies, the key characteristics necessary for enhanced replication of macaque-tropic HIV-1 clones remains poorly understood.

We have constructed PTM-tropic HIV-1 viruses (HSIV-vif) by replacing the *vif* genes with *vif* from highly pathogenic PTM-adapted SIVmne027 (Kimata et al., 1998; Kimata et al., 1999). These cloned viruses (CXCR4-tropic HSIV-vif_NL4-3_ and CCR5-tropic HSIV-vif_AD8_ and HSIV-vif_Yu2_) replicated better than their respective parental clones in PTM peripheral blood mononuclear cells (PBMCs) (Thippeshappa et al., 2011). Intravenous (IV) inoculation of PTMs with HSIV-vif_NL4-3_ showed low viral replication during the post-acute stages of infection through 44 wpi and small rebounds in viral titer at 64 and 72 wpi in juvenile PTMs (Thippeshappa et al., 2011). Furthermore, we observed that unlike pathogenic SIVmne, HSIV-vif_NL4-3_ replication is suppressed by type I interferon (IFN) treatment in PTM CD4^+^ T cells, perhaps suggesting that the IFN response during acute infection may limit virus replication in PTMs. Interestingly, we found that HSIV-vif_Yu2_ was resistant to IFNα-treatment in PTM CD4^+^ T cells in vitro, which may be due to envelope-mediated counteraction of IFNα-induced restrictions at the entry step of the viral life cycle (Thippeshappa et al., 2013). To further define important mutations in HSIV-vif that confer increased viral fitness PTMs, we isolated and characterized variant virus isolates from peripheral blood CD4^+^ T cells after 196-200 weeks post-inoculation (wpi) with HSIV-vif_NL4-3_ infection when there was CD4^+^ T cell depletion, and then performed a serial in vivo passaging experiment using a mixture of these virus isolates and different clones of HSIV-vif to define genetic characteristics associated with increased fitness and persistence.

## Methods

### Cell lines

TZM-bl cells were obtained from the NIH HIV Reagent Program (Derdeyn et al., 2000; Wei et al., 2002). TZM-bl and 293T cells were cultured in Dulbecco’s modified Eagle’s medium (DMEM) supplemented with 10% heat-inactivated fetal bovine serum (HI-FBS), 2 mM glutamine, 100 U of penicillin per ml and 100 μg of streptomycin per ml (p/s) (DMEM complete). The immortalized PTM CD4^+^ T-cells, obtained from Dr. Hans-Peter Kiem (Fred Hutchinson Cancer Research Center), were maintained in Iscove’s Modified Dulbecco’s Medium (IMDM) containing 10% HI-FBS, 2 mM glutamine, p/s, and 100 U/mL human interleukin 2 (IL-2) (Roche) (Munoz et al., 2009). CEMx174 were obtained from the American Type Culture Collection and cultured in RPMI with 10% HI-FBS, 2 mM glutamine, 100 U of penicillin per ml and 100 μg of streptomycin per ml (RPMI complete).

### Isolation of biological clones of HSIV-vif_NL4-3_

Total CD4^+^ T cells were isolated from 1 x 10^7^ peripheral blood mononuclear cells (PBMCs) recovered at 196 and 200 weeks post-infection of pigtailed macaque M08009 by negative selection using the Miltenyi nonhuman primate CD4^+^ T cell isolation kit (Miltenyi Biotech). The cells were isolated according to the manufacturer’s protocol. The M08009 CD4^+^ T cells were cocultured with the human T cell-B cell hybrid cell line, CEMx174 for up to 16 days. Supernatants after 7 and 16 days were assayed from HIV-1 p24 antigen by ELISA (Advanced Bioscience Laboratories). If positive, supernatants were passed through 0.45 μm syringe filters, aliquoted, and frozen at −80°C. The infectious titers of the stocks was determined by limiting dilution infection analysis using TZM-bl reporter cells and luciferase assay as described (Misra et al., 2018). Infectious virus recovered from CD4^+^ T cells at 196 wpi and 200 wpi were named C/196 and C/200, respectively.

### Coreceptor usage of HSIV-vif biological isolates

TZMbl cells (1 x 10^4^ cells per well) were plated in wells of a 96 well plate in DMEM complete with 30 μg/ml DEAE-dextran. In triplicate cultures, cells were treated with either the CXCR4 inhibitor, AMD3100, or CCR5 inhibitor, Maravoric, such that after adding 250 infectious units of C/196, C/200, or control viruses HSIV-vif_NL4-3_ or HSIV-vif_AD8_ the final concentrations of inhibitors were 1 μM, 500 nM, or 250 nM, and the final concentration of DEAE-dextran was 20 μg/ml. After two days of infection, the cells were washed once with PBS and lysed with Promega Glo Lysis buffer. Lysates were assayed for luciferase activity using the Promega luciferase assay system and tube luminometer according to the manufacturer’s instructions (Promega).

### Serum neutralization

Neutralizing antibody titers in serum specimens from PTMs infected with HSIV-vif_NL4-3_ were determined using a TZMbl-based neutralization assay as described previously (Wu et al., 2006). Serum samples were heat-inactivated at 56°C for 30 minutes prior to use.

### Plasmids

Construction of the HSIV-vif clones based on NL4-3, NL-AD8, and Bru-Yu2 has been reported before (Thippeshappa et al., 2011). To generate Vpr^+^ HSIV-vif_NL4-3_ and HSIV-vif_AD8_ clones, SphI to SalI fragment of HSIV-vif_NL4-3_ encompassing HIV *gag, pol*, SIV *vif* and HIV-1 vpr genes was cloned into pCR2.1 TOPO vector. SIV *vpx* start codon and two additional ATG codons upstream of the HIV-1 *vpr* start codon were mutated by Quickchange mutagenesis (Stratagene) and the sequence between the stop codon of *vif* and start codon of *vpr* were deleted. After mutagenesis, SphI and SalI fragment was cloned back into HSIV-vif_NL4-3_ and HSIV-vif_AD8_. Similarly, SphI to SalI fragment of HSIV-vif_Yu2_ was cloned into pCR2.1 TOPO vector, ATG codons upstream of *vpr* were mutated, and cloned back into HSIV-vif_Yu2_ (**Supplementary Fig. 1**). HSIV-vif-Vpx_Yu2_ clone was generated by cloning SacI to NcoI fragment of SIVmne027 *vpx* gene into pCR2.1 TOPO vector containing SphI to SalI fragment of HSIV-vif_Yu2_. SphI to SalI fragment containing full length *vpx* was cloned back into HSIV-vif_Yu2_ (**Supplementary Fig. 2**).

**Figure 1:**
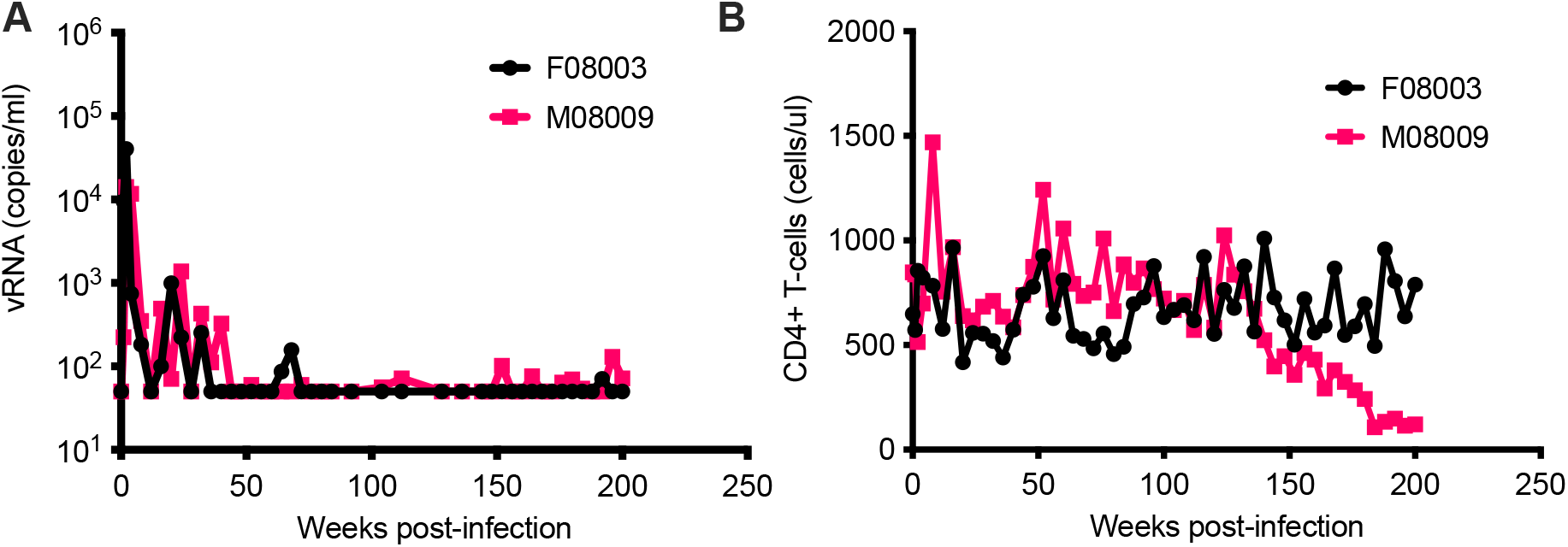
long-term monitoring of HSIV-vif_NL4-3_ infected PTMs. Two juvenile PTMs (F08003 and M08009) were inoculated intravenously with HSIV-vif_NL4-3_. Plasma viral RNA loads (A) and CD4^+^ T-cell counts (B) were measured at various time points post-infection. Data up to 90 wpi has been published previously (Thippeshappa et al., 2011). PTM M08009 shows gradual decline in CD4^+^ T cells at late-stage of infection.

### Virus stocks of HSIV-vif molecular clones for infection

Virus stocks were generated by transfection of 293T cells with each plasmid clone of HSIV-vif using Fugene 6 or X-tremeGENE 9 DNA transfection reagent according to the manufacturer’s protocol (Roche). Infectious titers were determined by limiting dilution infection analysis using TZM-bl indicator cells and the amount of virus in supernatants was measured by HIV-1 p24^gag^ antigen ELISA (Advanced Bioscience Laboratories).

### Western blot

293T cells, seeded the day before, in 6 well plates were transfected with HSIV-vif plasmids using Fugene 6 or X-tremeGENE 9 DNA transfection reagent (Roche/Sigma). At 48 hour post-transfection, cell culture supernatants were used to concentrate viral particles by centrifugation at 23600 x g for 1 hour at 4°C. Viral particles were mixed with 2X SDS samples buffer and separated by SDS-PAGE using Tris-HCl ready gels (Bio-Rad). Cell lysates were prepared as previously described (Thippeshappa et al., 2011). Proteins were transferred to either nitrocellulose or PVDF membrane and probed with rabbit antiserum to Vpr, Nef, or Vpx. Goat anti-rabbit IgG HRP (Promega) was used as secondary antibody. Antiserum to Vpr (Catalog # ARP-11836), Nef (Catalog # ARP-2949), Vpx (Catalog # ARP-2609), and anti-HIV-1 p24 Gag monoclonal (ARP-6458) were obtained from NIH HIV reagent program.

### Replication in PTM CD4 T cell line, PTM PBMCs, and human monocyte-derived macrophages

For viral replication assays, PTM PBMCs and immortalized PTM CD4^+^ T-cells were infected as previously described (Thippeshappa et al., 2011; Thippeshappa et al., 2013). PTM PBMCs were isolated using Ficoll hypaque gradient method. PBMCs were activated with concavalin A (7μg/ml) for 3 days. Cells were then washed twice and resuspended in RPMI complete media containing 40 U/ml IL-2 (Sigma) and cultured for 2 days. Approximately 1 x 10^6^ activated PBMCs were infected at a multiplicity of infection (MOI) of 0.01. To compare viral replication in PTM CD4^+^ T-cells, approximately 2.5 x 10^5^ or 5 x 10^5^ cells in 100 U/ml IL-2 containing IMDM were infected with viruses at a MOI of 0.01 or 0.05. Human MDMs were generated using previously described methods (Biesinger et al., 2010) and infected with the Vpr^+^ and Vpr-HSIV-vif. After three hours of incubation, the cells were washed twice with phosphate buffered saline (PBS) or complete medium to remove unbound virus. Infected cells were then resuspended in RPMI complete media containing IL-2. To study the effect of IFNa, 200 U/ml of Interferon-aA/D (IFN-aA/D or IFNa, Sigma) was added to the culture media. To monitor replication of HSIV-vif clones, supernatants were harvested every 2-4 days for measurement of HIV-1 p24^gag^ antigen using ELISA kit (Advanced Bioscience Laboratories or ExpressBio).

### Serial in vivo passaging in PTMs

Four PTMs specific pathogen free for simian T lymphotropic virus type 1, SIV, simian retrovirus type D, and herpes B virus were enrolled for the study. All animals were housed and cared for in accordance with the guidelines of the American Association for Accreditation of Laboratory Animal Care and the Animal Care and Use Committee of the University of Washington. In passage 1: two PTMs were inoculated intravenously (IV) with a mixture of CXCR4 and CCR5-tropic viruses. In passage 2: pooled peripheral blood from passage 1 PTMs collected at 14 wpi was used for inoculation of 1 PTM. In passage 3, week 8 peripheral blood from passage 2 PTM was used for transfusion into an additional PTM. At several time points post-inoculation, peripheral blood was drawn for CD4^+^ T-cell count determinations and isolation of plasma, sera, and PBMC.

### Plasma viral loads, CD4 T cell counts, and antibody response

Plasma viral load measurements were determined by using the Roche Amplicor HIV-1 monitor test, version 1.5 according to the manufacturer’s protocol. CD4^+^ T-cell counts were determined as previously described (Polacino et al., 2007). HIV-1-specific antibodies were measured by ELISA as previously described, using gradient-purified and disrupted whole HIV-1 virions as the capture antigen (Hu et al., 1989; Polacino et al., 2007).

### Cloning of IMCs by long-range PCR

The following steps were performed to generate IMCs (**Supplementary Fig. 3**).

#### Nested PCR amplification of near full length genome (NFLG)

Proviral DNA was isolated from 1×10^6^ PBMCs using Zymo Reseach Quick-DNA miniprep kit. In the first round PCR, 1 to 2μl of proviral DNA (approximately 50 to 100ng) in a 25μl reaction was amplified using the following primers: FWD-1: AAATCTCTAGCAGTGGCGCCCGAACAG and REV-1: TGAGGGATCTCTAGTTACCAGAGTC. Reaction mix contained 1× High Fidelity Buffer, 2 mM MgSO_4_, 0.2 mM dNTPs, and 0.025 U/μL Platinum Taq High Fidelity (Invitrogen). PCR conditions for the first round were 94°C for 2 min, then 94°C for 30 s, 64°C for 30 s, and 68°C for 10 min for 3 cycles; 94°C for 30 s, 61°C for 30 s, and 68°C for 10 min for 3 cycles; 94°C for 30 s, 59°C for 30 s, and 68°C for 10 min for 3 cycles; 94°C for 30 s, 57°C for 30 s, and 68°C for 10 min for 21 cycles; and then 68°C for 10 min. 1 μl of first-round PCR reaction product was amplified using following primers FWD-2: ACAGGGACTTGAAAGCGAAAG and REV-2: CTAGTTACCAGAGTCACACAACAGACG. Reaction mix and PCR conditions were identical to the first round PCR. PCR products were visualized on 1% agarose gel and gel eluted using QIAquick gel extraction kit.

#### Vector PCR

10ng of HSIV-vif_NL4-3_ was amplified with following primers: 3LTR-V-F90: TGTGTGACTCTGGTAACTAGAGATCCCTCAGACCCTTTTAGTCAGTGTGGAAAATCTCTAGCACCCAGGAGGTAGAGGTTGCAGTGAGC and 5HIV-R2: CTTTCGCTTTCAAGTCCCTGTTCGGGCGCCA in a 50μl reaction volume. Platinum superfi II high fidelity DNA polymerase (Thermofisher) was used for amplification of vector PCR product. Vector PCR product was visualized on 1% agarose gel and gel eluted using QIAqucik gel extraction kit.

#### NEBuilder HiFi DNA assembly

NFLG PCR product and vector PCR product were mixed with a minimum of 1:5 ratio in a 20μl reaction volume containing 10μl of HiFi assembly mix. After 1 hour of incubation at 50’c, 5μl of reaction mix was used for transformation of NEB STABLE cells. Miniprep plasmids were isolated using QIAprep spin miniprep kit. Plasmids containing full-length genomes were screened by restriction enzyme digestion with SalI and BamHI enzymes.

To determine if the plasmids containing full length clones generate infectious virus, transfection supernatants were generated by transfecting 293T cells with proviral clones. Infectious nature of the supernatants was determined by infecting TZM-bl cells.

### GenBank accession numbers

Sequences of HSIV-P3-114, HSIV-P3-161, and HSIV-P3-284 are deposited under accession numbers MZ146778, MZ146779, and MZ146780 respectively.

## Results

### Isolates of HSIV-vif_NL4-3_ from late stage infection evolved resistance to host immune responses

We previously reported HSIV-vif_NL4-3_ could persistently infect PTMs (Thippeshappa et al., 2011). We continued to monitor the viral loads in two of those HSIV-vif_NL4-3_-infected PTMs for nearly 4 years (**Fig. 1A**). Although viral RNA was below the detection limit at most of the late time points measured, viral DNA could be detected in PBMCs through 200 weeks post-inoculation (wpi). Additionally, we recovered infectious virus from PBMCs at 196 wpi and 200 wpi, suggesting that the virus had been replicating in the animals for nearly 4 years. Interestingly, one of the animals, despite low or undetectable viral loads, showed a gradual decline and then stably depressed CD4^+^ T cell counts, suggesting disease progression (**Fig. 1B**).

We recovered infectious virus by coculturing peripheral blood CD4^+^ T cells from infected PTM (M08009, **Fig. 1**) at 196 wpi and 200 wpi with CEMx174 cells (biological isolates C/196 and C/200). Partial genome sequencing analysis of PCR amplified fragments of C/196 and C/200 indicated that both isolates were clonal (**Supplementary Fig. 5**). C/196 and C/200 were susceptible to inhibition by AMD3100 but not Maravoric, suggesting that viral isolates were only CXCR4-tropic (**Supplemental Fig. 4**). We next determined the replication capacity of C/196 and C/200 in PTM CD4^+^ T cells (**Fig. 2**). C/196 replicated to higher levels in PTM CD4^+^ T cells than the parental clone, HSIV-vif_NL4-3_. Although it remained susceptible to IFNa, its peak replication in the presence of IFNa was higher than that of the parental virus (**Fig 2A and 2B**). While C/200 displayed more limited replication, it was not affected by the addition of IFNa (**Fig. 2C**). We also observed that both C/196 and C/200 were relatively neutralization resistant to sera from M08009, the animal in which it evolved, as well as sera from the other PTM, F08003, that had been infected with HSIV-vif_NL4-3_ (**Table 1**). The emergence of immune escape variants of HSIV-vif_NL4-3_ suggest that it had persistently replicated in PTMs.

**Figure 2:**
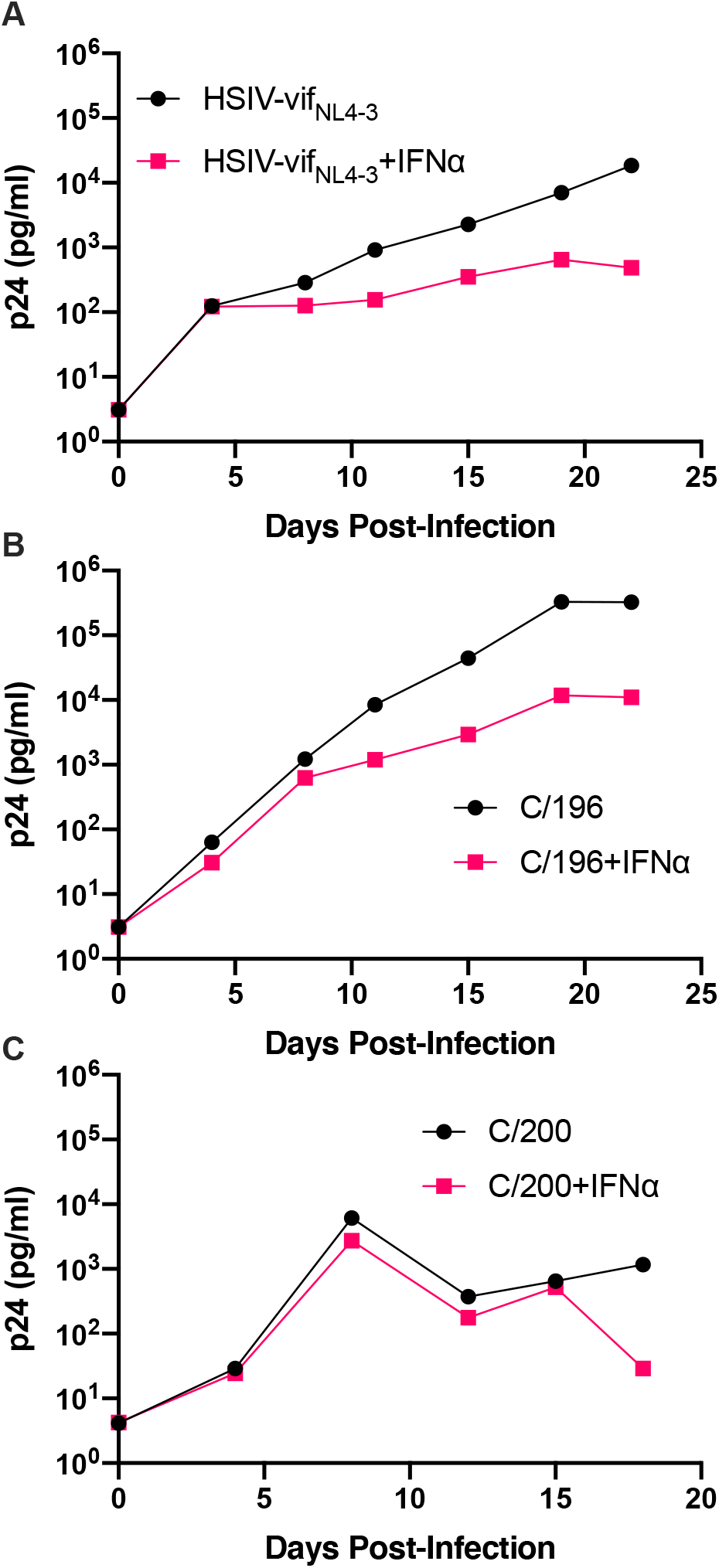
Replication kinetics of biological isolates of HSIV-vif_NL4-3_ (C/196 and C/200). PTM CD4+ T cells were infected in duplicate at a MOI of 0.01 with the parental HSIV-vif_NL4-3_ (A) or variant isolates C/196 (B) and C/200 (C) in the presence or absence of IFNa (200U/ml) in the culture media. Virus supernatants were collected every 3 to 4 dpi and p24 was quantified by ELISA.

**Table 1:**
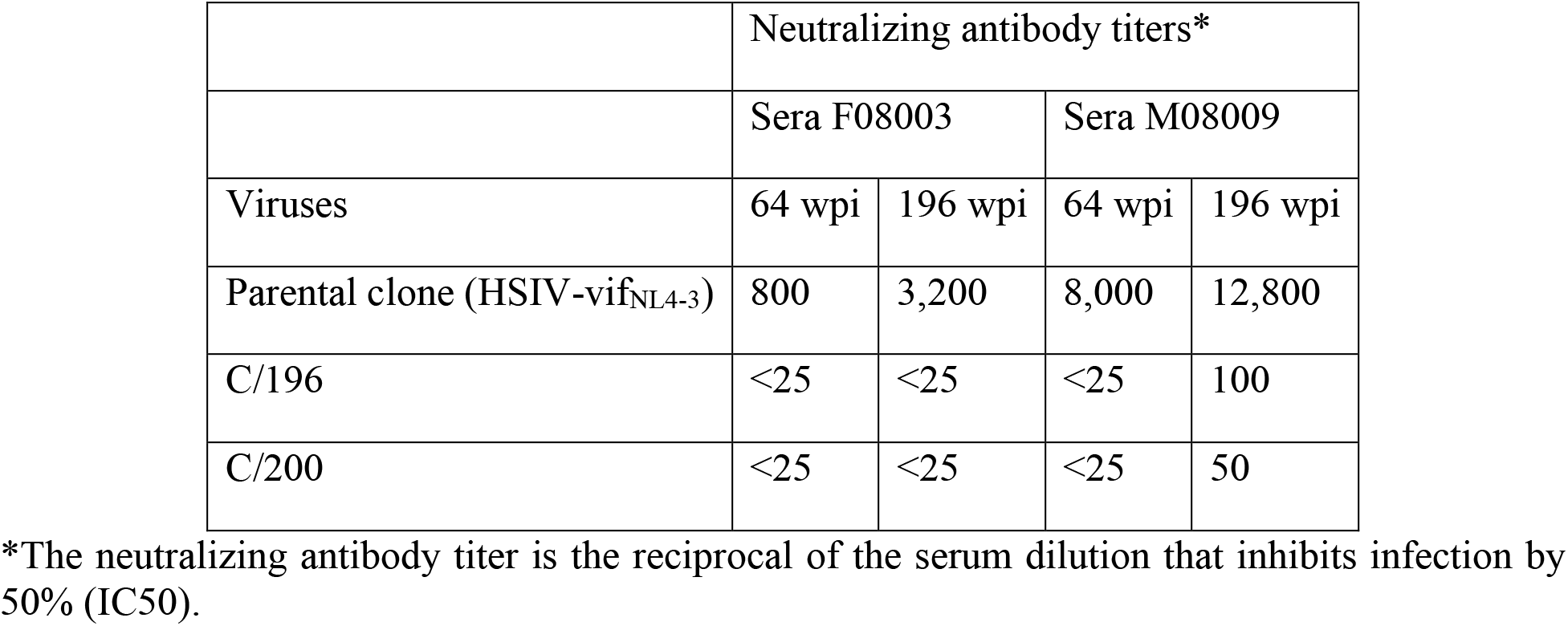
Serum neutralizing antibody titer against wild type versus late-isolates of HSIV-vif_NL4-3_

### Characterization of HSIV-vif clones expressing Vpr and Vpx

An explanation for the low but persistent replication of HSIV-vif_NL4-3_ in vivo is that the virus is attenuated because it does not express accessory proteins necessary for more robust viral replication. Two possible proteins that could enhance replication of HSIV-vif_NL4-3_ are the HIV-1 Vpr or SIV Vpx. Since we introduced SIV *vif*, which includes a partial open reading frame (ORF) for *vpx*, into HIV-1 backbone, we determined if it had affected the expression of HIV-1 *vpr*. Indeed, the HIV-1 Vpr protein was not observed in virions of HSIV-vif clones, HSIV-vif_NL4-3_ or HSIV-vif_AD8_ (**Fig. 3A**). We deteremine that this is because singly spiced HIV-1 *vpr* RNA from HSIV-vif is generated using the splice acceptor site within the SIV *vif* gene (**Supplementary Fig. 1**). The transcript, therefore, includes the partial ORF for the SIV Vpx protein upstream of the translational initiation site for HIV-1 Vpr, which may interfere with its expression (**Supplementary Fig. 1**).

**Figure 3:**
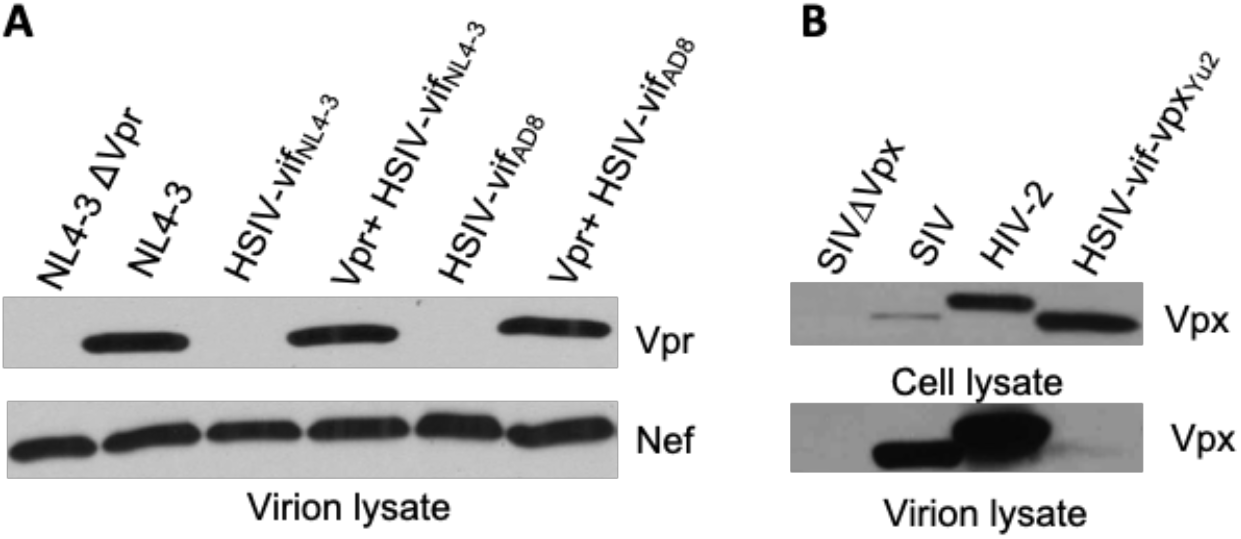
Vpr and Vpx expression from HSIV clones. 293T cells were transfected with HSIV clones. At 48 hours post-transfection, virus supernatants were collected and concentrated by centrifugation. Cell and virion lysates were analyzed by western blot using antibody to HIV-1 Vpr Vpx, and Nef.

Vpr is a small 96 amino acid (14 kDa) protein that is not required for HIV-1 replication *in vitro*. However, it is conserved among all primate lentiviruses, indicating its importance for pathogenesis (Tristem et al., 1992; Tristem et al., 1998). Therefore, we modified HSIV-vif clones to express Vpr. We deleted the sequence between the stop codon of *vif* and start codon of *vpr* and disrupted the *vpx* translational start codon and other ATG codons upstream of the *vpr* initiation site by site directed mutagenesis (**Supplementary Fig. 1 & 7**). Mutation of these 3 ATG codons, one of which results in M181L in SIV Vif, resulted in expression of Vpr, which is incorporated into progeny virions (Vpr^+^ HSIV-vif_NL4-3_ and Vpr^+^ HSIV-vif_AD8_, **Fig. 3A**). Additionally, since Vpx counteracts the function of SAMHD1 (Hrecka et al., 2011; Laguette et al., 2011) and is essential for replication of SIV in macaques (Belshan et al., 2012; Shingai et al., 2015) and because HSIV-vif already has a partial ORF for vpx, we also generated an HSIV-vif derivative carrying the full-length *vpx* gene **Supplementary Fig. 2**). We used HSIV-vif_Yu2_, which is IFN-resistant (Thippeshappa et al., 2013), to generate HSIV-vif-vpx carrying the full-length *vpx* gene (named HSIV-vif-vpx_Yu2_). By Western blot using rabbit anti-serum to HIV-2_Rod_ Vpx protein, Vpx could be detected in the cell lysates of SIV, HIV-2, and HSIV-vif-vpx_Yu2_, but not SIVΔVpx. However, it was not detected in virion lysates of HSIV-vif-vpx_Yu2_ (**Fig. 3B**) as HIV-1 does not have determinants in p6 Gag required for virion incorporation of Vpx (Sunseri et al., 2011).

We tested the effect of Vpr and Vpx expression on HSIV-vif replication in an immortalized PTM CD4^+^ T-cell line (Munoz et al., 2009). PTM CD4^+^ T-cells were infected with Vpr^-^ (HSIV-vif_NL4-3_, HSIV-vif_AD8_ and HSIV-vif_Yu2_), Vpr^+^ (Vpr^+^HSIV-vif_NL4-3_, Vpr^+^HSIV-vif_AD8_ and Vpr^+^HSIV-vif_Yu2_), HSIV-vif-vpx_Yu2_, or wild type HIV-1 (NL4-3, NL-AD8 and Bru-Yu2) viruses at an MOI of 0.01. Vpr^+^ HSIV-vif viruses, and HSIV-vif-vpx_Yu2_ replicated in PTM CD4^+^ T-cells to similar levels as Vpr^-^ viruses (**Fig. 4**). Expectedly, wild type HIV-1 (NL-AD8 and Bru-Yu2) failed to replicate (**Fig. 4**).

**Figure 4:**
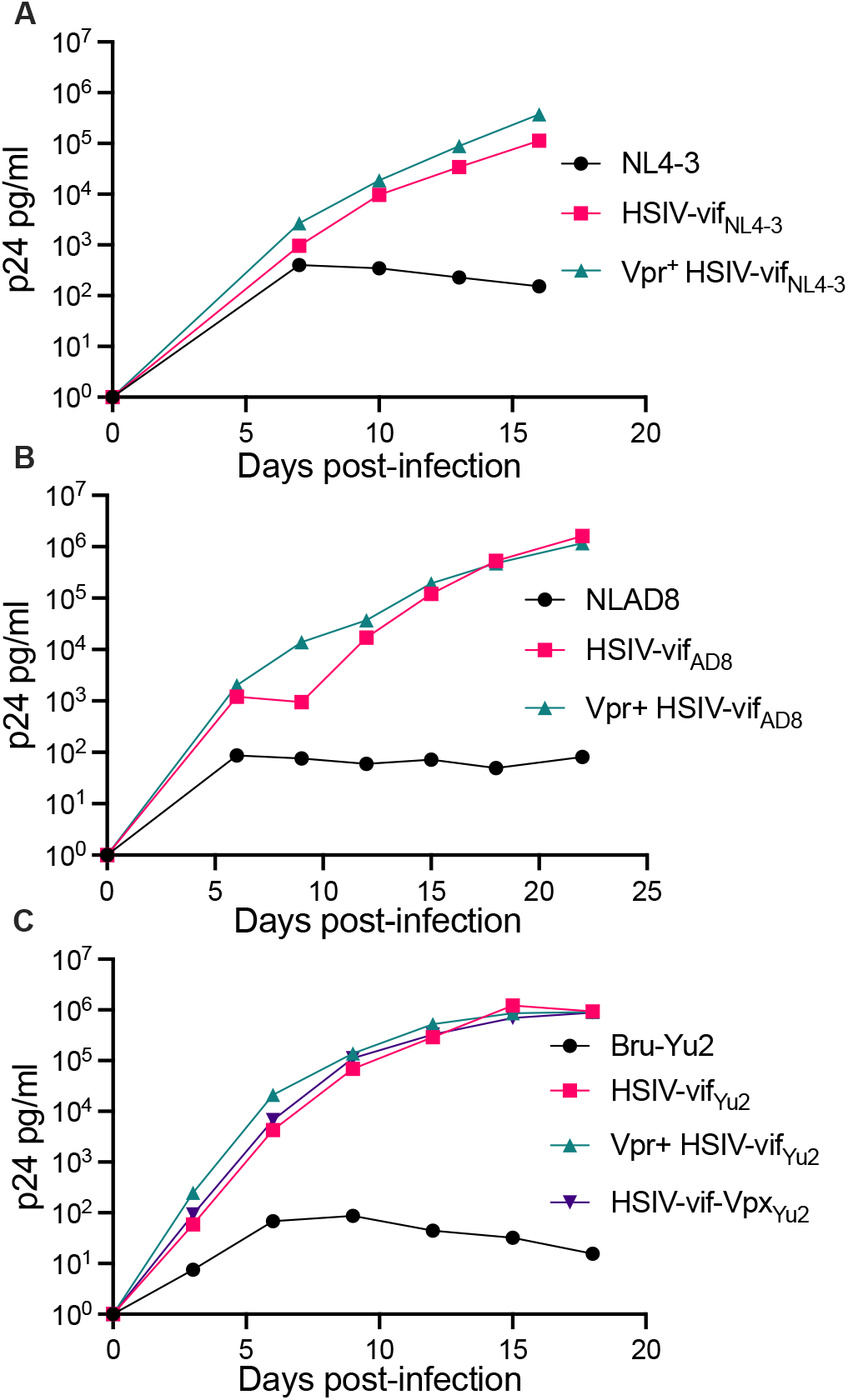
Vpr and Vpx expressing HSIV replicate in PTM CD4 T cells. Cells were infected in duplicate with HSIV-vif_NL4-3_ (A), HSIV-vif_AD8_ (B), and HSIV-vif_Yu2_ (C) variants at an MOI of 0.01. Virus supernatants were collected at 3 to 4 days post-infection and assayed for p24 levels.

We also determined the replication capacity of *vpr* and *vpx* carrying HSIV clones in human monocyte-derived macrophages (MDMs). Human MDMs were generated using previously described methods (Biesinger et al., 2010) and infected with the Vpx^+^, Vpr^+^ and Vpr-HSIV-vif clones at an MOI of 0.01. Vpr^+^ HSIV-vif_AD8_ (**Fig. 5A**)and Vpr^+^ HSIV-vif_Yu2_ (**Fig. 5B**) replicated to similar levels as Vpr^-^ HSIV-vif_AD8_ and Vpr^+^ HSIV-vif_Yu2_, but slightly less than wild type HIV-1 NL-AD8 and Bru-Yu2 (**Fig. 5A&B**). Additionally, HSIV-vif-vpx_Yu2_ replicated as well as the parental HIV-1 Bru-Yu2 (**Fig 5B**).

**Figure 5:**
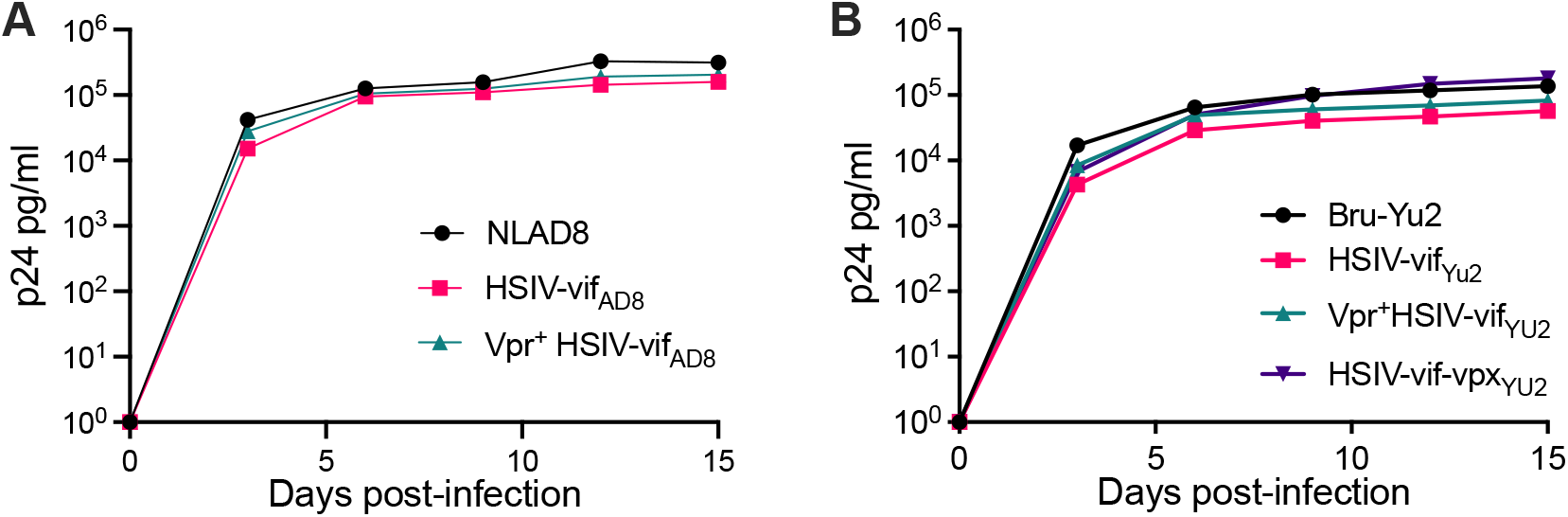
Vpr and Vpx expressing HSIV replicate in MDMs. Cells were infected in duplicate with HSIV-vif_AD8_ (A) and HSIV-vif_Yu2_ (B) clones expressing Vpr or Vpx or neither at an MOI of 0.01. Virus supernatants were collected at 3 to 4 days post-infection and assayed for p24 levels.

### Serial in vivo passaging of HSIV-vif

Because HSIV-vif_NL4-3_ replication in the initial PTM experiment was low with peak viremia <10^5^ copies/ml (**Fig. 1**) (Thippeshappa et al., 2011), we conducted animal to animal transfer of infected PTM peripheral blood to adapt HSIV-vif to PTMs. For this experiment, the initial inoculum contained a mixture of CXCR4-(C/196 and C/200) and CCR5-tropic viruses (Vpr^+^ HSIV-vif_AD8_, Vpr^+^ HSIV-vif_Yu2_, and HSIV-vif-vpx_Yu2_). At 14 wpi, pooled blood from infected PTMs (Z09080 and Z09067) was used to inoculate a naïve macaque (Z13086). At 8 wpi, peripheral blood from Z13086 was transferred into an additional PTM (Z13098). Interestingly, all the macaques showed a peak viremia close to or above 10^5^ copies/ml and the viral loads persisted for at least 20 wpi (**Fig. 6A**). Furthermore, increases in antibody titer over time suggest that all PTMs were persistently infected with HSIV-vif (**Fig. 6B**). However, CD4^+^ T cell decline was not observed in the infected PTMs (**Fig. 6C**).

**Figure 6:**
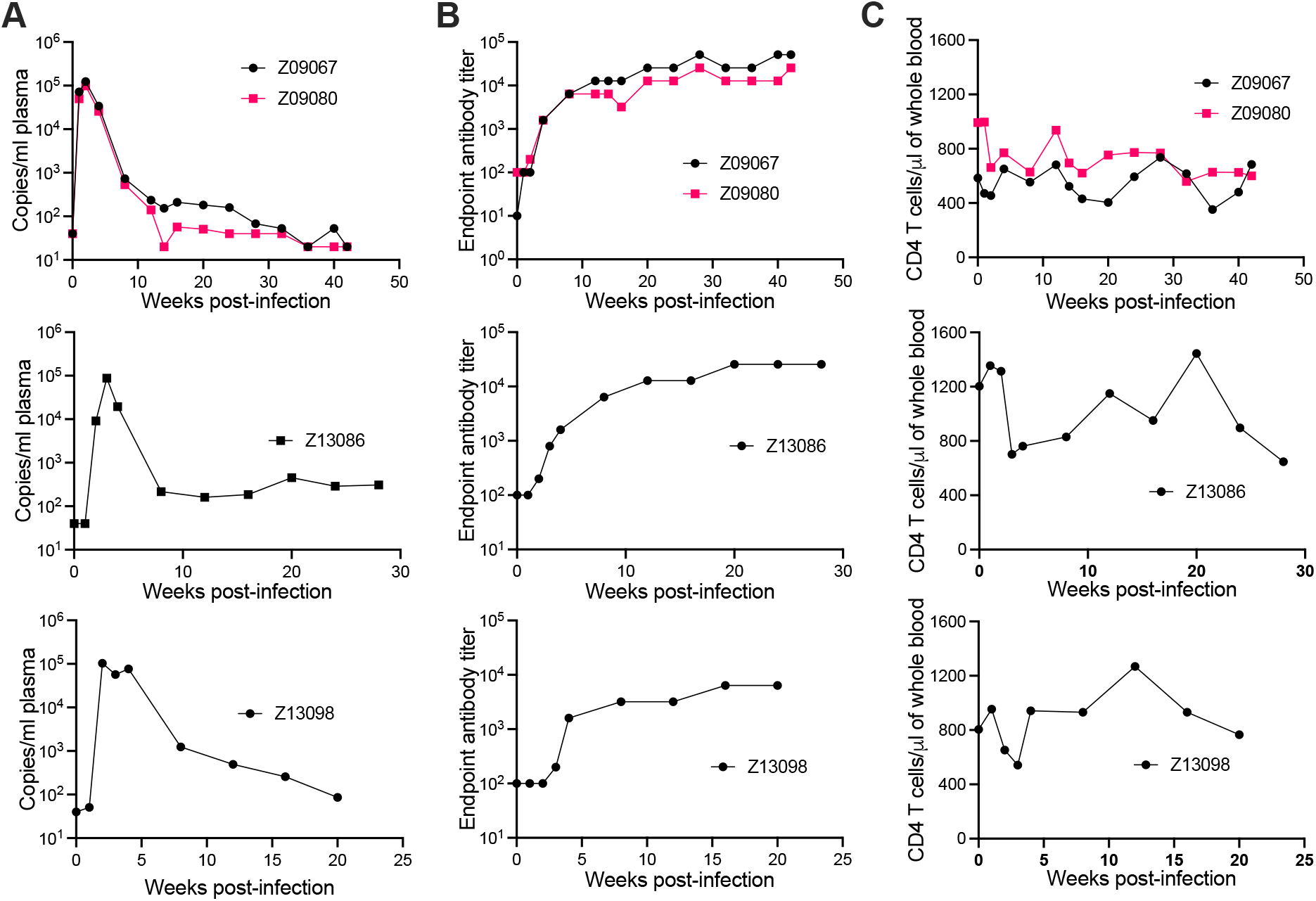
Passaging of HSIV to generate pathogenic variants. Passage 1 macaques were infected with a mixture of viruses. At 14 wpi, pooled blood from infected PTMs (Z09080 and Z09067) was used to inoculate a naïve macaque (Z13086) and then week 8 blood from Z13086 was passaged through an additional PTM (Z13098). At several time points post-inoculation, peripheral blood was drawn for measuring plasma viral loads (A), antibody titer (B), and CD4 T cell counts (C).

### Cloning and characterization of HSIV IMCs

We generated IMCs from proviral DNA isolated from PBMCs of the passage 3 macaque (Z13098). Near full-length proviral genomes (NFLG) were amplified using nested PCR as described by Heiner et al (Hiener et al., 2017) with slight modifications in PCR conditions. NFLGs was cloned into a vector PCR product containing 5’ LTR sequences (amplified from HSIV-vif_NL4-3_ plasmid) using NEBuilder HiFi assembly mix (**Supplementary Fig. 3**). Briefly, ends of NFLG PCR product and vector PCR product containing 5’LTR sequences overlap with each other, which can be assembled using NEBuilder HiFi assembly mix. We screened nearly 300 colonies to identify 54 plasmids with full length genomes. To determine if they could produce infectious virus, full length clone plasmids were transfected into 293T cells to generate virus. Infectious nature of the supernatants was determined by infecting TZM-bl cells. Three of the 54 plasmid clones tested (HSIV-P3-114, HSIV-P3-161, and HSIV-P3-284) produced measurable infectious virus. DNA sequencing showed that the 3 IMCs were closely related to the C/196 and C/200 biological clones of HSIV-vif_NL4-3_ (**Table 2, Supplementary Fig. 5 and 6**). Importantly, the three IMCs had mutations throughout the genome suggestive of adaptation to PTMs (**Table 2**). Most of the mutations in Env and Nef were seen in the biological clones of HSIV-vif_NL4-3_ (C/196 and C/200) recovered from PTM M08009 (**Table 2**), suggesting that these mutations have persisted through 3 additional in vivo passages. In the Vpr^+^ HSIV-vif clones, we had mutated SIV *vpx* ATG codon to ACG (silent mutation) and ATG codon at amino acid position 181 in SIV *vif* to TTG, which codes for leucine (M181L substitution) (**Supplementary Fig 1**). However, SIV *vpx* start codon was present in the recovered IMCs. Additionally, ATA codon was present at amino acid position 181, which codes for isoleucine (M181I substitution). Further, all the recovered IMCs had the deletion of bases between SIV *vif* stop codon and *vpr* start codon (**Supplementary Fig 7**), suggesting that these clones could express Vpr protein. We confirmed the expression of Vpr from HSIV-P3 IMCs by western blot using rabbit anti-sera against Vpr protein (**Fig 7**). Interestingly, virion associated Vpr was higher than that for the VPR^+^ HSIV-vif_NL4-3_ clone. The recovery of Vpr^+^ HSIV-vif clones similar to C/196 and C/200 suggests recombination occurred between a Vpr^+^ HSIV-vif clone and a biological isolate of HSIV-vif_NL4-3_.

**Figure 7:**
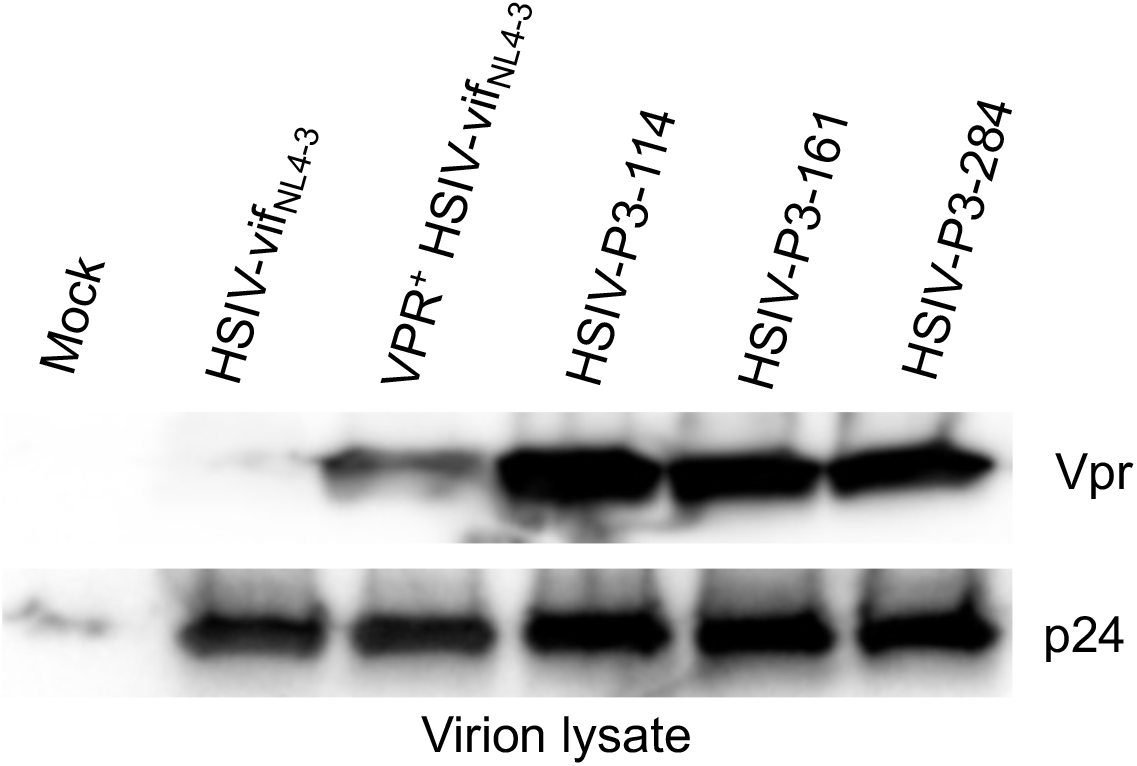
Vpr expression from HSIV-P3 IMCs clones. 293T cells were transfected with HSIV-P3 IMCs. At 48 hours post-transfection, virus supernatants were collected and concentrated by centrifugation. virion lysates were analyzed by western blot using antibody to HIV-1 Vpr and p24.

**Table 2:**
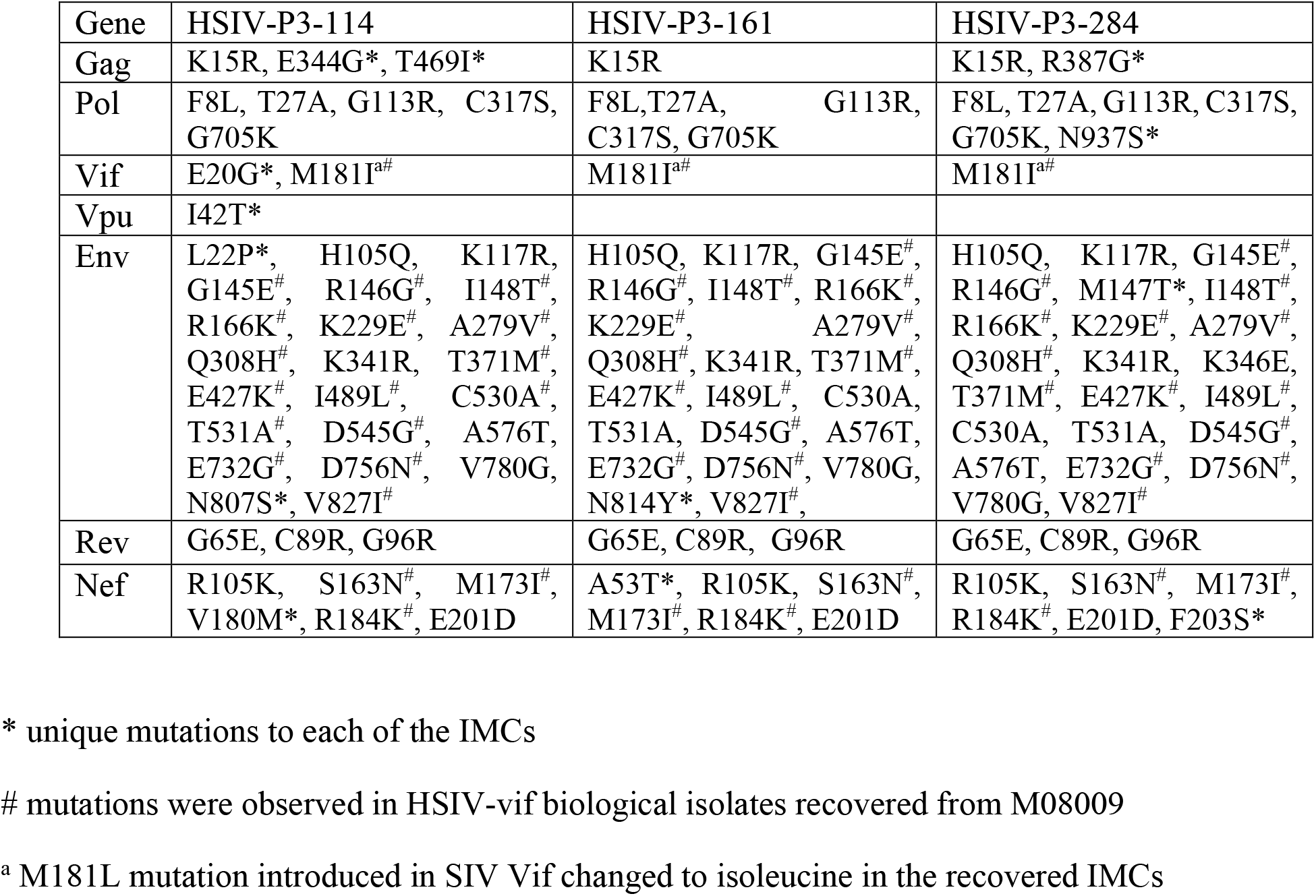
Non-synonymous mutations observed in HSIV-P3 IMCs

We determined if HSIV-P3 IMCs (HSIV-P3-114, HSIV-P3-161, and HSIV-P3-284) replicate in PTM PBMCs. PBMCs isolated from different donor PTMs were activated with concavalin A for 3 days and maintained in IL-2 containing media for 2 days. Activated PBMCs were infected with HSIV-P3 IMCs and Vpr^+^ HSIV-vif_NL4-3_ at an MOI 0.01. Viral supernatants were collected at various days post-infection to assay for p24 levels. We observed that HSIV-P3 IMCs replicated with different efficiency in PBMCs from different donor PTMs (**Fig. 8**). Among the three HSIV-P3 IMCs, HSIV-P3-284 replicated to the highest level in activated PBMCs.

**Figure 8:**
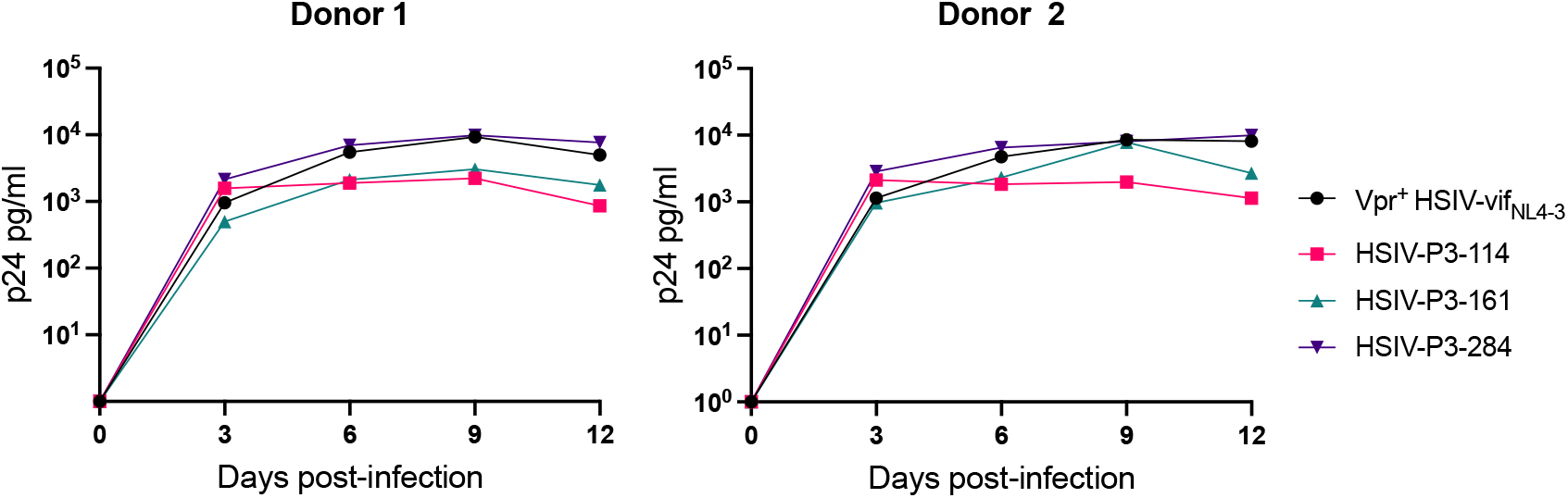
HSIV-P3 IMCs replicate in PTM PBMCs. PBMCs from different donor PTMs (Z14117 and Z14112) were activated with concavalin A for 3 days and maintained in media containing IL-2 (40U/ml) for 2 days. Cells were then infected in duplicate with HSIV-P3 IMCs or parental Vpr^+^ HSIV-vif_NL4-3_ virus at an MOI of 0.01 and supernatants were assayed for p24 by ELISA every 3 to 4 dpi.

## Discussion

Since first generation SHIV constructs replicated poorly in macaques, serial in vivo passages were conducted to enhance their infectivity or replicative capacity (Luciw et al., 1995; Joag et al., 1996; Reimann et al., 1996; Igarashi et al., 1999; Chen et al., 2000; Song et al., 2006). We previously reported the construction of HSIV-vif_NL4-3_ and replication trend in juvenile and newborn PTMs (Thippeshappa et al., 2011). Although HSIVvif_NL4-3_ persisted for nearly 4 years, the peak viremia was below 10^5^ copies/ml, rapidly declined, and was intermittently detectable thereafter in juvenile PTMs. Therefore, we conducted serial in vivo passaging of HSIV-vif in PTMs using a mixture of different molecular clones and variants as an initial inoculum. We report that peak plasma viral RNA levels were greater than 1×10^5^ viral RNA copies/ml and continuously detectable for 20-30 wpi. To further characterize HSIV-vif selected in PTMs, we PCR cloned and generated IMCs from the passage 3 macaque using DNA from week 4 PBMCs from passage 3 PTM. Our characterization indicated that the selected variants appear to be Vpr expressing recombinants of the HSIV-vif_NL4-3_ biological isolates from the long-term infected PTM with depressed CD4^+^ T cell counts. These data suggest that neutralization resistant variants which had evolved in association with CD4^+^ T cell decline and acquired the ability to express the HIV-1 Vpr had a fitness advantage for replication in PTMs.

We have previously reported potential reasons for the attenuated replication of HSIV-vif_NL4-3_ in PTMs. In those studies, we noticed that HSIV-vif_NL4-3_ does not degrade PTM APOBEC3 family restriction factors as well as highly pathogenic SIVmne027 (Thippeshappa et al., 2011). We also observed that replication of HSIV-vif_NL4-3_ is inhibited in the presence of IFNa in PTM CD4 T cells (Thippeshappa et al., 2013). This perhaps suggests that HSIV-vif_NL4-3_ may not be able to overcome type I IFN response induced during acute stage of infection. Here, we also show that HSIV-vif clones do not express Vpr due to interference by the partial *vpx* open reading frame (ORF) in the singly spliced *vpr* mRNA. We speculate that absence of Vpr expression may have affected viral replication of HSIV-vif_NL4-3_ in the initial inoculation of PTMs (Fig. 1).

HIV-1 Vpr incorporates into virions through an interaction with p6 of Gag (Bachand et al., 1999; Selig et al., 1999). In vitro studies have attributed several biological function to Vpr, which include: i) cell cycle arrest and apoptosis (Di Marzio et al., 1995; He et al., 1995; Jowett et al., 1995; Planelles et al., 1995; Re et al., 1995; Bartz et al., 1996; Goh et al., 1998; Stewart et al., 1999; Stewart et al., 2000; Zhang and Bieniasz, 2020); ii) nuclear import of viral DNA (Popov et al., 1998a; Popov et al., 1998b; Le Rouzic et al., 2002; Riviere et al., 2010); iii) regulation of viral gene expression (Forget et al., 1998; Subbramanian et al., 1998; Vanitharani et al., 2001; Yurkovetskiy et al., 2018); iv) infection of nondividing cells (Balliet et al., 1994; Connor et al., 1995; Campbell and Hirsch, 1997; Miller et al., 2017); v) modulation of immune responses (Ayyavoo et al., 2002; Muthumani et al., 2004; Majumder et al., 2005; Muthumani et al., 2005; Majumder et al., 2008; Okumura et al., 2008; Doehle et al., 2009; Khan et al., 2020); and vi) interaction with uracil DNA glycosylase (UNG2), a DNA repair enzyme that specifically removes uracil from DNA, and reduction of G to A mutations during reverse transcription (Mansky et al., 2000; Chen et al., 2004; Ahn et al., 2010). Since Vpr performs multiple functions during HIV replication, we hypothesized that absence of HIV-1 Vpr expression in HSIV-vif_NL4-3_ may affect persistent viral replication in PTMs.

It is interesting that even without Vpr expression, HSIV-vif_NL4-3_ persisted for nearly 4 years (Fig 1). While the functions of Vpr have not been clearly defined *in vivo*, the potential importance of Vpr for HSIV-vif replication in pigtails is supported by pathogenesis studies of SIVmac, which demonstrate deletion of either Vpr or Vpx alone or together attenuates viral replication and ability to cause disease (Lang et al., 1993; Gibbs et al., 1995). Interestingly, Vpr null viruses reverted to Vpr expressing virus in SIV-infected macaques (Lang et al., 1993; Hoch et al., 1995), suggesting the importance of Vpr for in vivo pathogenesis. We speculated that the HIV-1 Vpr expressing HSIV may replicate better than the parental Vpr-HSIV-vif_NL4-3_ in vivo. Therefore, we generated HSIV derivatives expressing HIV-1 Vpr by introducing mutations in ATG codons upstream of the Vpr start codon. Since Vpx performs similar roles as Vpr and SIV *vif* gene already has partial open reading frame for *vpx*, we also generated a HSIV-vif derivative expressing the full length *vpx* gene. We cloned full length vpx gene into HSIV-vif_Yu2_ backbone, as we have previously shown that this clone resists IFN treatment in PTM CD4 T-cells. HSIV-vif clones expressing either the HIV-1 *vpr* or SIV *vpx* were replication competent *in vitro*. However, accessory proteins such as Vpr and Vpx are not necessary for HIV-1 or SIV replication *in vitro*. Therefore, it is difficult to show the impact of Vpr or Vpx expression for HSIV-vif replication using *in vitro* studies. Infecting PTMs with different clones would be a better method to define the significance of the HIV-1 *vpr* for HSIV-vif replication. Indeed, in our passage studies, we used both HIV-1 Vpr- and Vpr^+^ HSIV to determine the importance of HIV-1 Vpr for in vivo pathogenesis in PTMs. Recovery of Vpr^+^ HSIV IMCs from the passage 3 macaque again suggests a role for Vpr in in vivo pathogenesis.

In our studies to characterize persistent HSIV-vif variants, we have developed and standardized a rapid and robust approach to generate IMCs from proviral DNA. We screened 54 plasmids for their ability to generate infectious virus. Out of which, only 3 generated infectious virus, which roughly correspond to 5% of total clones. This is not surprising as 90 to 95% of the proviral DNA is non-infectious (Ho et al., 2013; Bruner et al., 2016; Hiener et al., 2017). Interestingly, recovered IMCs from passage 3 macaque were Vpr^+^ HSIV-vif_NL4-3_. This suggests a possible recombination between Vpr^+^ HSIV-vif clones (either HSIV-vif_AD8_ or HSIV-vif_Yu2_) with biological isolates C/196 and C/200 recovered virus from M08009. Two observations suggest a recombination event: First, the 3 recovered IMCs had deletion of bases between SIV *vif* stop codon and HIV-1 *vpr* start codon. Second, most mutations observed in *env* and *nef* were already present in the HSIV-vif_NL4-3_ biological clones (C/196 and C/200) from M08009. Therefore, the recombination event to generate Vpr^+^ HSIV indicates the importance of the HIV-1 Vpr for HSIV-vif pathogenesis in vivo.

Although we used mixture of CXCR4- and CCR5-tropic viruses for inoculation into passage 1 PTMs, it is interesting that CXCR4-tropic HSIV-vif_NL4-3_ persisted through 3 passages. We have previously reported that SIV variants emerging during late-stage disease have a higher replicative capacity and increased pathogenicity (Kimata et al., 1999). Similar observations have also been made with SHIV-1157ipd (Song et al., 2006). Therefore, recovered virus (C/196 or C/200) isolated during late stage in our study may have greater fitness for replication in PTMs. Since recovered virus was also neutralization resistant, it may have helped the virus overcome antibody responses during additional passages. We also observed several non-synonymous mutations throughout the genome of HSIV-P3-IMCs. Most of the mutations were shared among the three HSIV-P3 IMCs. However, HSIV-P3 IMCs also had mutations unique to each of the clones. We speculate that these mutations could help the virus overcome restriction factors, better utilize host dependency factors, or they could help the virus escape adaptive immune responses.

In conclusions, our results suggest that serial in vivo passaging improves HSIV replication and persistence in PTMs. Identification of several nonsynonymous mutations in IMCs recovered from passage 3 macaque also indicated that serial in vivo passaging helps in acquisition of mutations.

Since these mutations are in the context of replication competent virus, they may play a significant role in the replication and pathogenesis in vivo. However, a shortcoming of the studies is the limited duration of the passage experiment and limited number of animals used for the study. While variants of HSIV-vif_NL4-3_ that acquired the ability to express the HIV-1 Vpr appear to have a selective advantage, the short duration of the experiments was insufficient to determine if the selected variants had increased pathogenicity. Further in vivo passaging of HSIV-P3 IMCs with longer follow-up periods will be necessary to verify their increased replication fitness, and to generate pathogenic variants with enhanced replication capacity. Development of such pathogenic variants will be valuable as challenge viruses for preclinical evaluation of novel vaccines and therapeutics, as these HSIV clones have all the HIV immunologic and vaccine targets such as Gag, Pol, Env, Tat, Rev, and Nef. Furthermore, establishment of HIV reservoirs in this model also provides an avenue for developing therapeutic vaccination approaches targeting HIV Gag, Pol, and Env, apart from testing latency reversal agents and cure strategies.

## Author contributions

RT: designed studies, performed experiments, wrote manuscript; PP: Coordinated and analyzed data from animal studies, and edited manuscript; SSC: designed and performed experiments, edited manuscript; KT and AM performed experiments and edited manuscript; SLH: obtained funding, coordinated animal studies and edited manuscript; DK; obtained funding, coordinated studies, and edited manuscript; JTK: obtained funding, designed studies, performed experiments, wrote manuscript.

## Acknowledgements

The following reagents were obtained through the NIH HIV reagent program, Division of AIDS, NIAID, NIH: Polyclonal Anti-Human Immunodeficiency Virus Type 2ROD Vpx Protein (antiserum, Rabbit), ARP-2609, contributed by Dr. Lee Ratner, Washington University; Polyclonal Anti-Human Immunodeficiency Virus Type 1 Vpr Protein, Residues 1 to 50 (antiserum, Rabbit), ARP-11836, contributed by Dr. Jeffrey Kopp; Polyclonal Anti-Human Immunodeficiency Virus Type 1 Nef Protein (antiserum, Rabbit), ARP-2949, contributed by Dr. Ronald Swanstrom; Anti-Human Immunodeficiency Virus 1 (HIV-1) p24 Gag Monoclonal (#24-3), ARP-6458, contributed by Dr. Michael Malim; Human Immunodeficiency Virus 1 (HIV-1), Strain NL4-3 Infectious Molecular Clone (pNL4-3), ARP-2852, contributed by Dr. M. Martin; Human Immunodeficiency Virus 1 (HIV-1) AD8 Infectious Molecular Clone (pNL(AD8)), ARP-11346, contributed by Dr. Eric O. Freed; Clone, ARP-1350, contributed by Dr. Beatrice Hahn and Dr. George M. Shaw; TZM-bl Cells, ARP-8129, contributed by Dr. John C. Kappes and Dr. Xiaoyun Wu.

## Supplementary Figure legends

**Supplementary Fig. 1:**
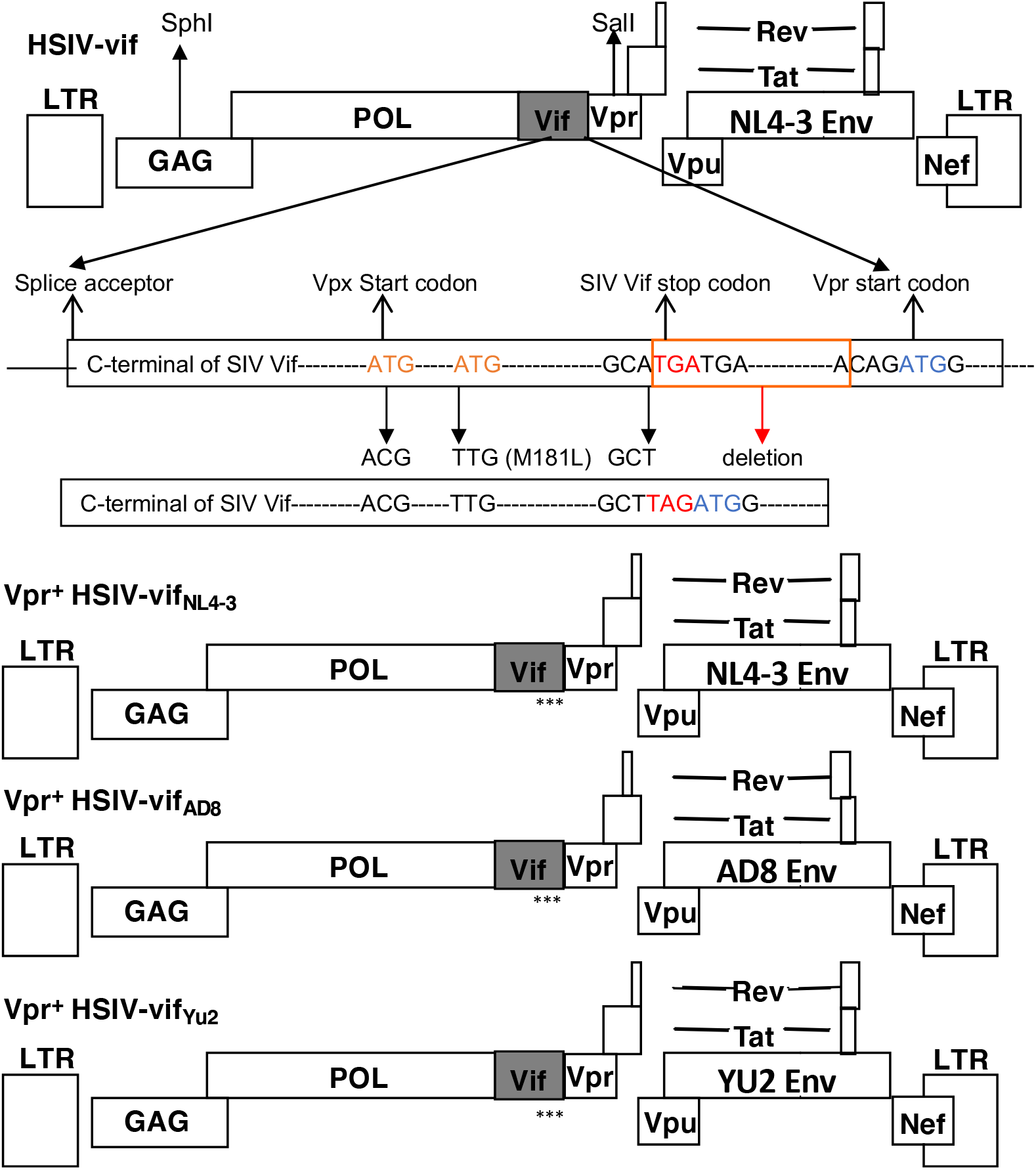
Construction of Vpr expressing HSIV-vif clones. SphI to SalI fragment of HSIV-vif_NL4-3_ encompassing HIV *gag, pol*, SIV *vif* and HIV-1 vpr genes was cloned into pCR2.1 TOPO vector. SIV *vpx* start codon and two additional ATG codons upstream of the HIV-1 *vpr* start codon were mutated by Quickchange mutagenesis and the sequence between the stop codon of *vif* and start codon of *vpr* were deleted. After mutagenesis, SphI and SalI fragment was cloned back into HSIV-vif_NL4-3_ and HSIV-vif_AD8_. Similarly, SphI to SalI fragment of HSIV-vif_Yu2_ was cloned into pCR2.1 TOPO vector, ATG codons upstream of *vpr* were mutated, and cloned back into HSIV-vif_Yu2_. *Approximate location of mutations introduced in the *vif* gene.

**Supplementary Fig. 2:**
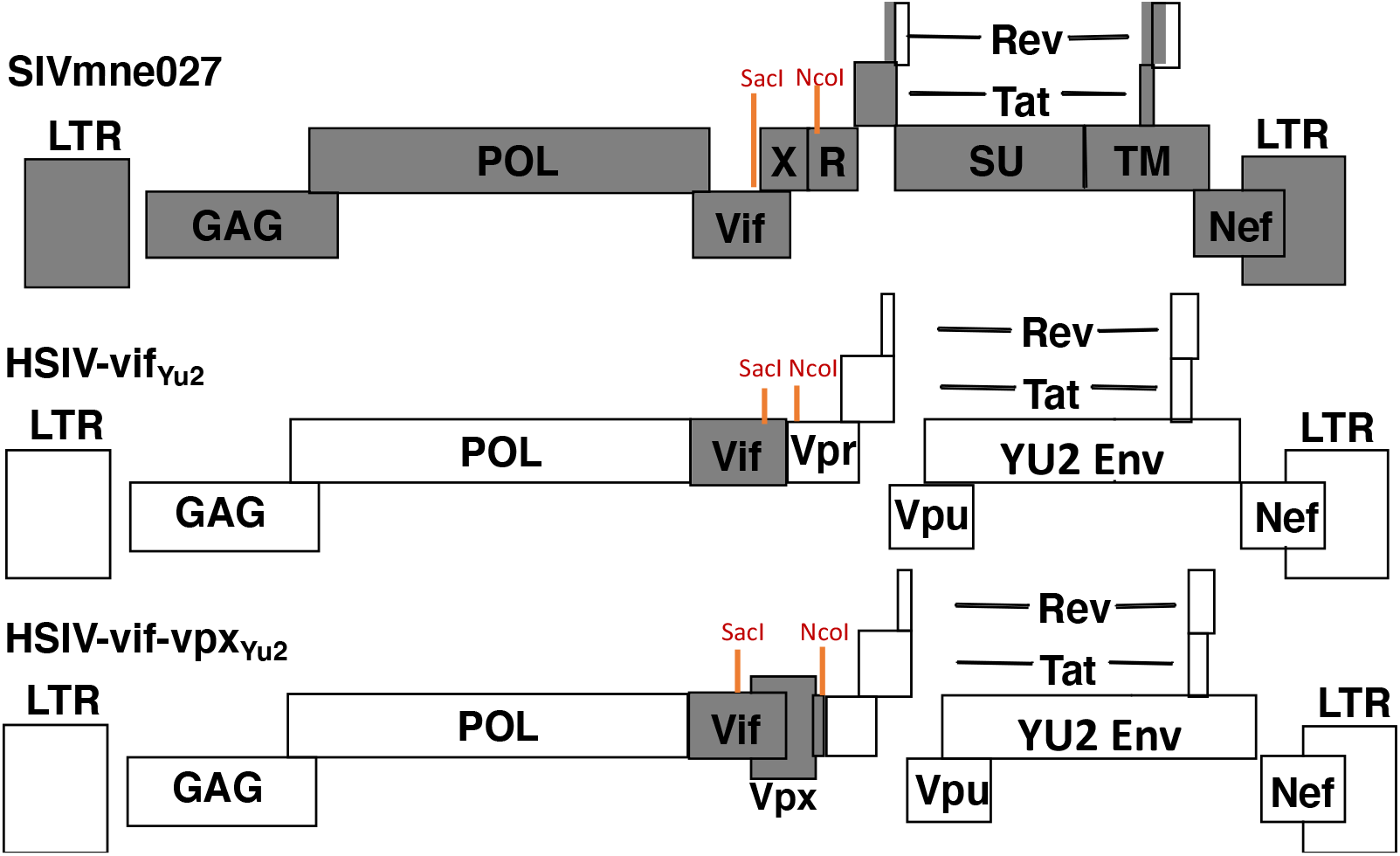
Construction of Vpx expressing HSIV-vif_Yu2_. SacI to NcoI region of HSIV-vif_Yu2_ was replaced with SacI to NcoI region of SIVmne027 to generate HSIV-vif-vpx_Yu2_.

**Supplementary Fig. 3:**
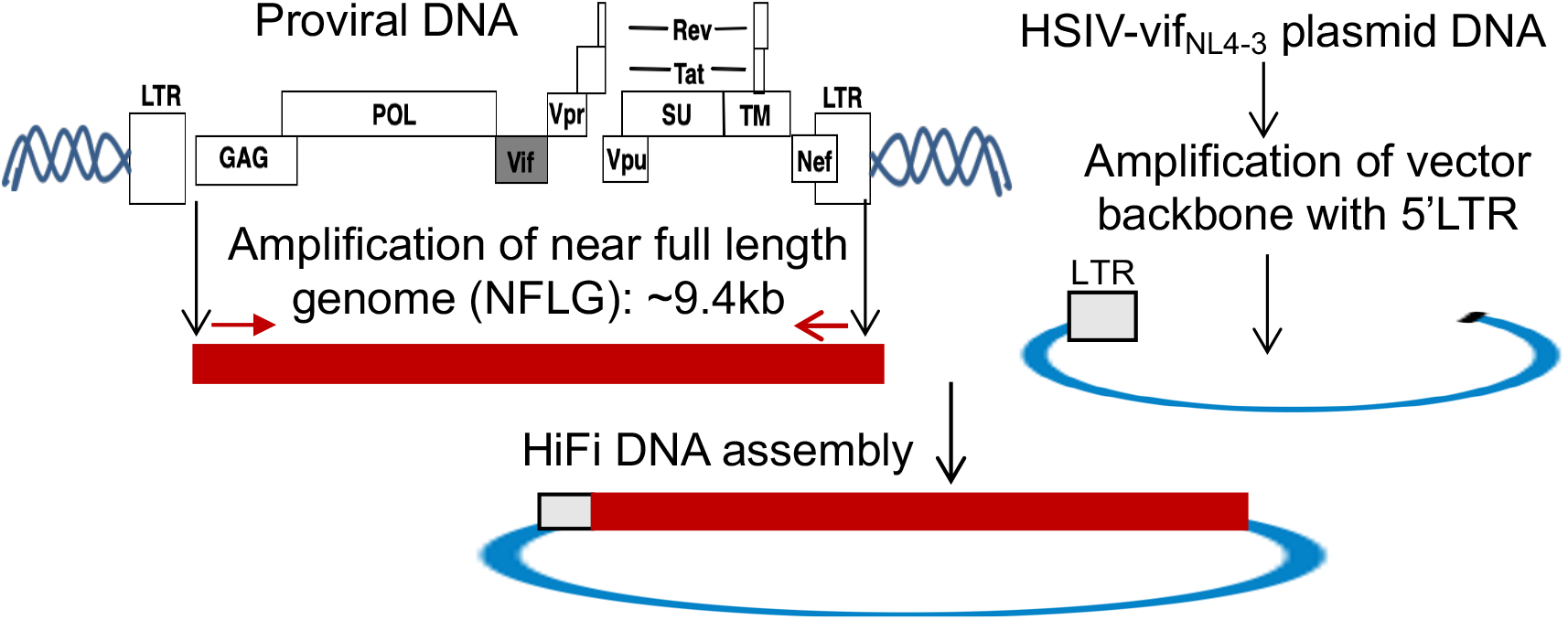
Schematic of HiFi DNA assembly approach to generate full length clones. Near full length genomes (NFLG) were amplified using nested PCR and cloned into a vector PCR product containing 5’ LTR sequences (amplified from HSIV-vif_NL4-3_ plasmid) using NEBuilder HiFi assembly mix.

**Supplementary Fig. 4:**
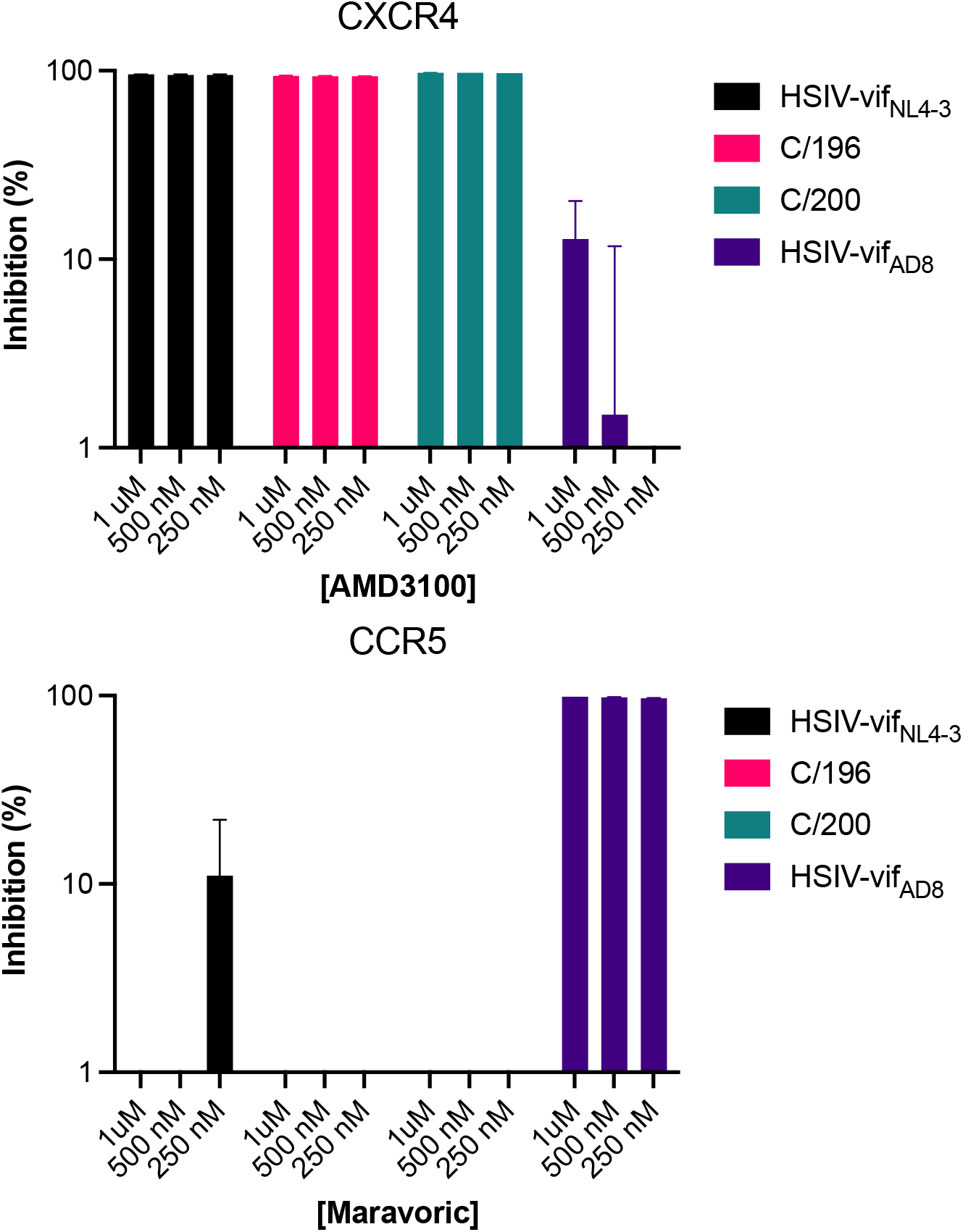
Coreceptor usage of biological clones C/196 and 200-2. TZM-bl cells were infected with different viruses in the presence of increasing concentrations of CXCR4 and CCR5 inhibitors. 48 hours later cell lysates were assayed for luciferase activity using plate based luminometer.

**Supplementary Fig. 5:**
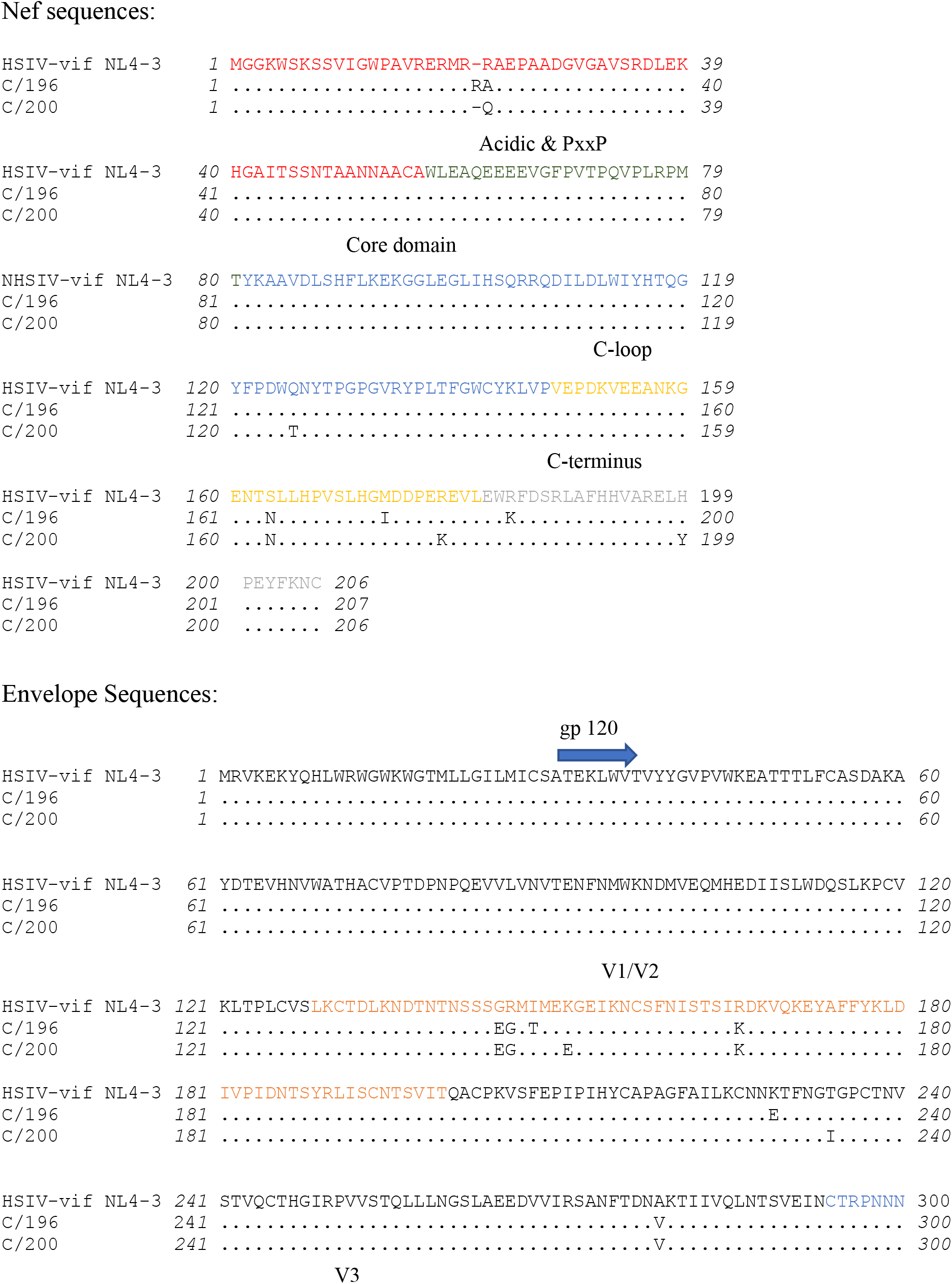
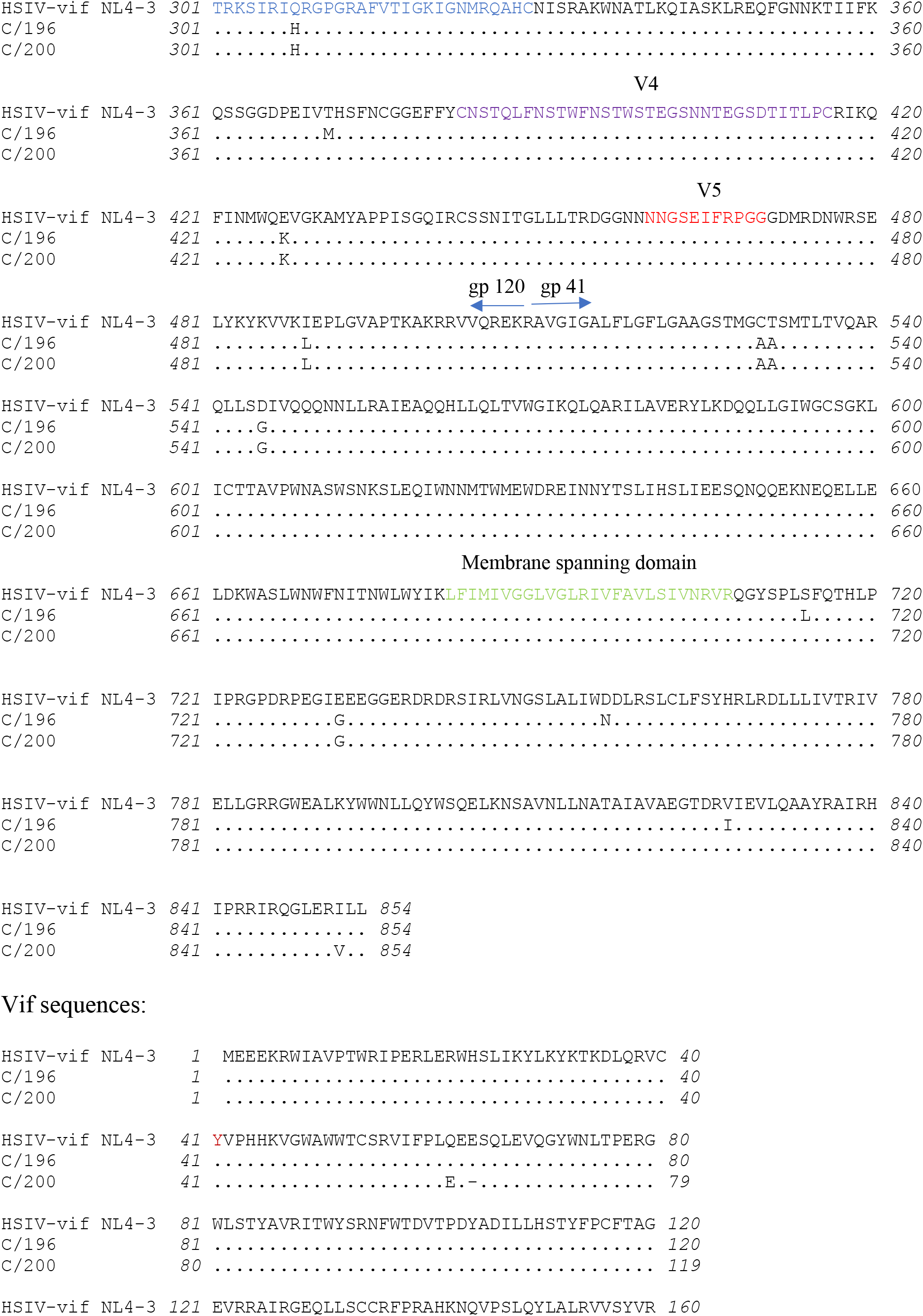
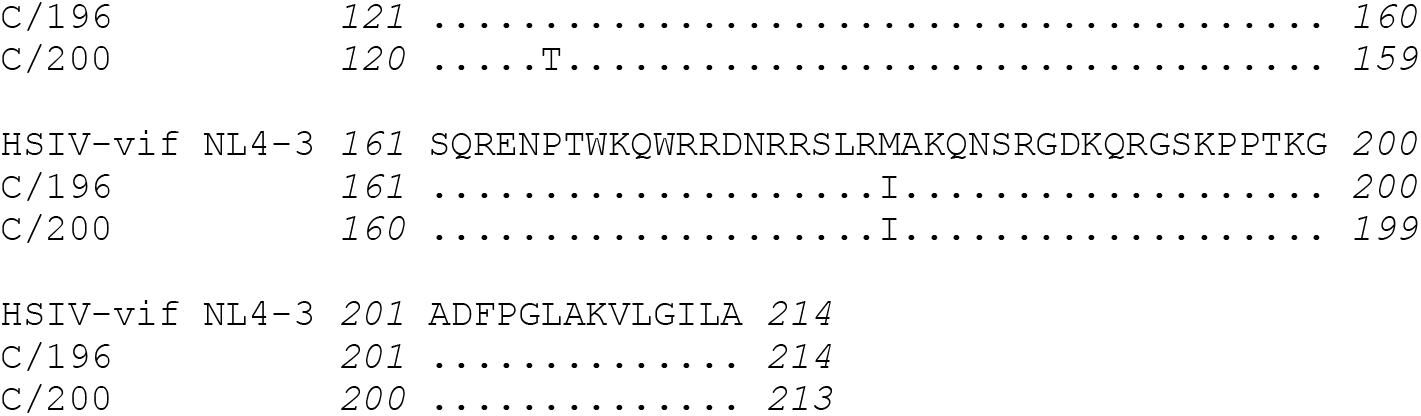
Sequence alignment of biological isolates (C/196 and C/200). Nef, Vif, and Envelope protein sequences of biological isolates recovered PTM MO8009 are aligned to parental HSIV-vif_NL4-3_ sequences.

**Supplementary Fig. 6:**
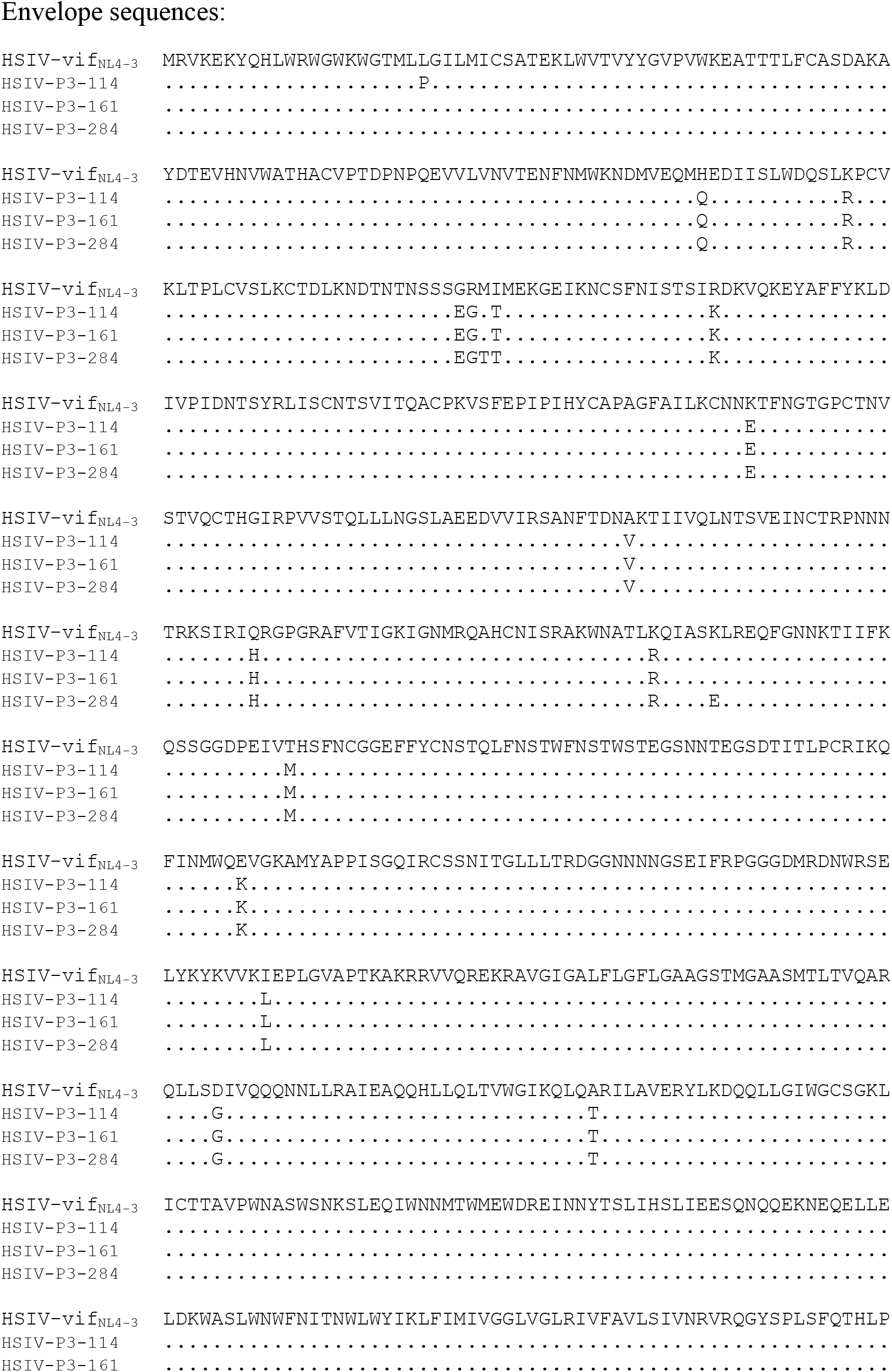
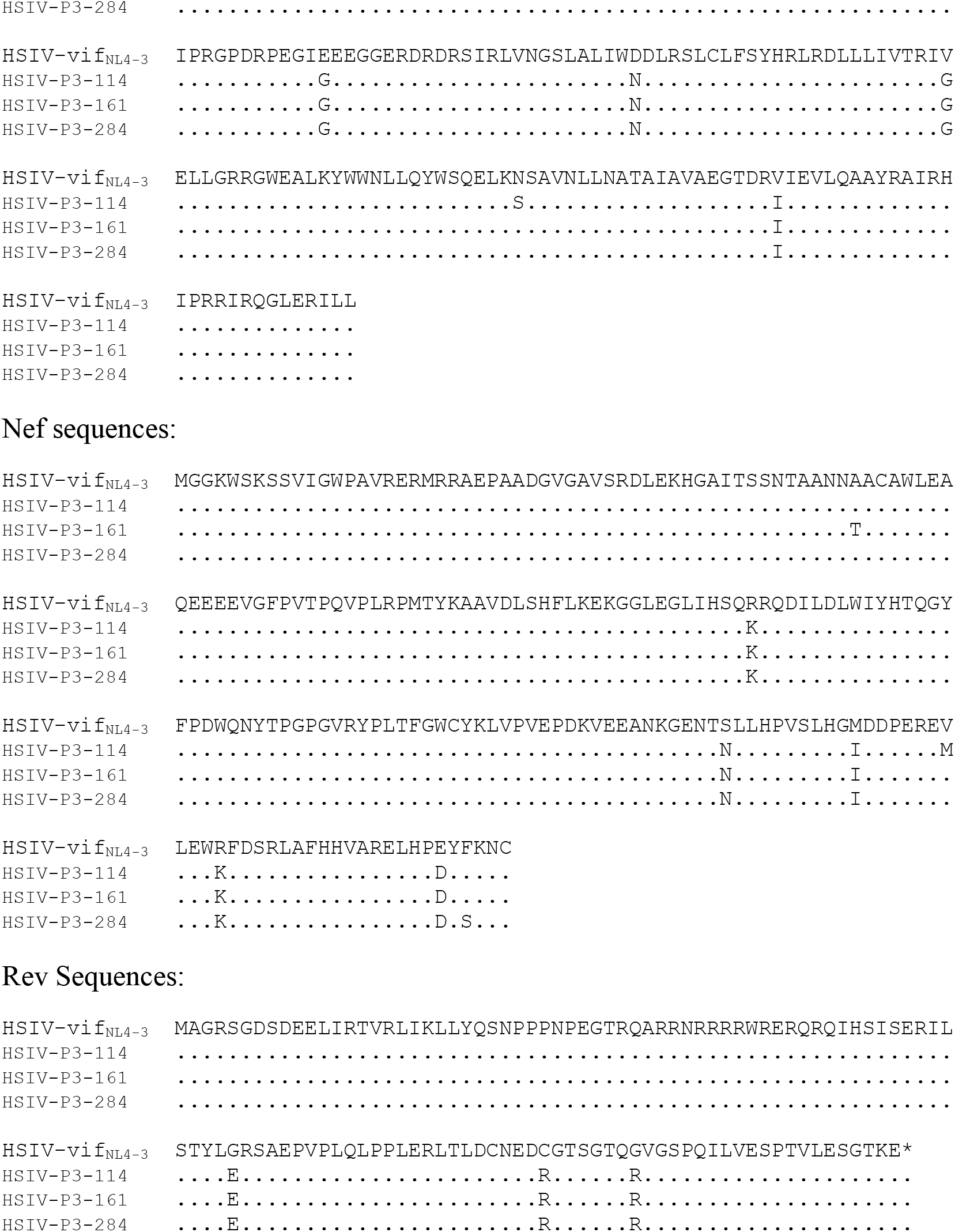
Sequence alignment of HSIV-P3 IMCs. Envelope, Nef, and Rev protein sequences of HSIV-P3 IMCs are aligned to parental HSIV-vif_NL4-3_ sequences.

**Supplementary Fig. 7:**
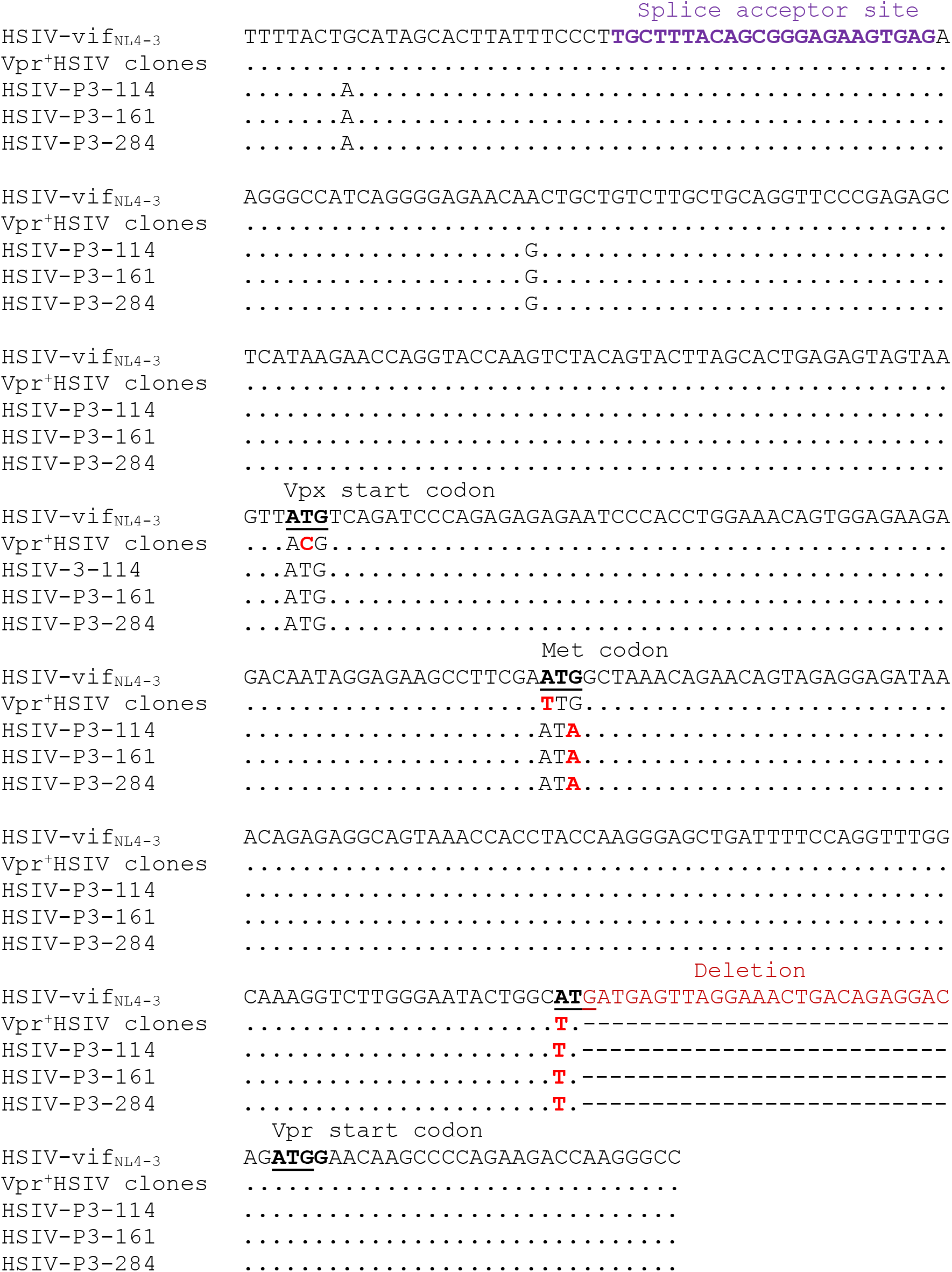
Sequence alignment of SIV vif and HIV-1 vpr region of HSIV-P3 IMCs with Vpr^+^ and Vpr-HSIV-vif_NL4-3_.

